# Optimized CART Cell Therapy for Metastatic Aggressive Thyroid Cancer

**DOI:** 10.1101/2024.03.15.585286

**Authors:** Claudia Manriquez Roman, Elizabeth L. Siegler, Justyna J. Gleba, Truc N. Huynh, Grace E. DeFranco, Aylin Alasonyalilar Demirer, Matthew L. Pawlush, Michael Redig, Skyeler M. Klinge, Long K. Mai, James L. Miller, Erin E. Miller, Brooke L. Kimball, Erin E. Tapper, R. Leo Sakemura, Carli M. Stewart, Ismail Can, Olivia L. Sirpilla, Jennifer M. Feigin, Kun Yun, Omar L. Gutierrez Ruiz, Hong Xia, Mehrdad Hefazi Torghabeh, Kendall J. Schick, Ekene J. Ogbodo, Gloria Olivier, Yushi Qiu, Robert C. Smallridge, Abba Zubair, Han W. Tun, John A. Copland, Saad S. Kenderian

**Affiliations:** Center for Regenerative Biotherapeutics, Mayo Clinic; Rochester, MN; T Cell Engineering Laboratory, Mayo Clinic; Rochester, MN; Division of Hematology, Mayo Clinic; Rochester, MN; Cancer Biology Department, Mayo Clinic; Jacksonville, FL; Department of Immunology, Mayo Clinic; Rochester, MN; Mayo Clinic Graduate School of Biomedical Sciences; Rochester, MN; Department of Molecular Pharmacology and Experimental Therapeutics, Mayo Clinic; Rochester, MN; Department of Biochemistry and Molecular Biology, Mayo Clinic; Rochester, MN; Department of Molecular Medicine, Mayo Clinic; Rochester, MN; Department of Business Development, Mayo Clinic; Rochester, MN; Division of Endocrinology, Internal Medicine Department, Mayo Clinic; Jacksonville, FL; Department of Laboratory Medicine and Pathology, Mayo Clinic; Jacksonville FL; Hematology/Oncology Division, Department of Medicine, Mayo Clinic; Jacksonville, FL

## Abstract

Most thyroid cancer deaths are attributed to a subset of poorly differentiated, metastatic tumors. To improve treatment options for aggressive thyroid cancers, we developed a novel thyroid-stimulating hormone receptor (TSHR)-targeted chimeric antigen receptor T (CART) cell therapy, which demonstrated antigen-specific activation and antitumor efficacy against TSHR^+^ cell lines *in vitro* and *in vivo*. However, de-differentiated thyroid cancers downregulate TSHR. We therefore developed a potent treatment strategy by combining our novel TSHR-CART cells with mitogen-activated protein kinase (MAPK) inhibitors, which redifferentiate thyroid tumors and upregulate TSHR expression. In patient-derived anaplastic thyroid cancer xenografts, combination therapy of TSHR-CART cells and MAPK inhibitors led to increased TSHR expression on the tumor tissue and significantly enhanced antitumor efficacy and prolonged survival compared to TSHR-CART monotherapy. Based on our data, we are launching a phase I clinical trial for TSHR-CART cell therapy alone or in combination with MAPK inhibitors in patients with metastatic thyroid cancers.

**STATEMENT OF SIGNIFICANCE:** Poor target selection and antigen escape limit CART cell efficacy in solid tumors. We developed TSHR-CART cells to treat differentiated thyroid cancers but observed TSHR downregulation in dedifferentiated thyroid cancers. We found that MAPK inhibitors restored TSHR expression and sensitized these cancers to TSHR-CART cell therapy.

## INTRODUCTION

Thyroid cancer is the most common endocrine cancer, and its incidence is rising in the US and globally.(*1, 2*) Thyroid cancer is projected to become the fourth leading type of cancer across the world, and the causes appear to be complex and multifactorial, including dietary changes and environmental exposures. Patients with thyroid cancer have widely different clinical outcomes depending on the pathological subtype and mutation profile. Metastatic differentiated thyroid cancer (DTC), poorly differentiated thyroid cancer (PDTC), and anaplastic thyroid cancer (ATC) account for a majority of thyroid cancer-related deaths, even though these subtypes only represent 2-3% of all thyroid cancer diagnoses.(*1, 3, 4*) Hurthle cell carcinoma (HCC, now termed oncocytic thyroid cancer(*5*)) is a subtype of DTC known to be aggressive, highly metastatic, and have lower rates of overall survival than follicular thyroid cancers. HCC is often resistant to radioiodine therapy and has no current FDA-approved treatment options.(*6, 7*) PDTC and ATC are aggressive and often refractory to radioiodine therapy and other standard therapies.(*8, 9*) SEER database analysis through 2015 reported a median overall survival of 4 months in patients with ATC, with disease-specific mortality of over 98%. A more recent analysis reported a median overall survival of 15.7 months in patients with ATC receiving newer targeted therapies.(*10*) Despite these advancements in treatment options, PDTC and ATC remain incredibly lethal.

Chimeric antigen receptor (CAR) T cell therapy has emerged as a potentially curative therapy in a subset of patients with hematological malignancies and has been FDA-approved for several B cell malignancies as well as for multiple myeloma.(*11–15*) With the advent of adoptively transferred engineered cellular therapies, there is a compelling rationale to apply CART cell therapy to treatment-resistant solid tumors. To date, the efficacy of CART cell therapy in solid tumors has been modest at best.(*16–21*) Due to a lack of tumor-specific targets for CART cell therapy in solid tumors, identifying an antigen that is both uniquely and universally expressed on cancer cells is critical for development of antigen-specific immunotherapy to avoid on-target off-tumor toxicities.(*20, 22, 23*)

Thyroid-stimulating hormone receptor (TSHR) is a surface glycoprotein receptor and a major regulator of thyroid function and metabolism. The TSHR mediates the activating mechanism of TSH to the thyroid gland, resulting in the growth and proliferation of thyrocytes as well as thyroid hormone production, including thyroxine (T4) and triiodothyronine (T3).(*24*) TSHR is predominantly expressed on the basolateral membrane of normal thyroid follicular cells.(*25*) TSHR expression is largely limited to the thyroid gland, with low levels of expression in additional tissues, including the thymus and testes (immune privileged) according to Human Protein Atlas and NIH GenBank databases. TSHR is abundantly expressed on thyroid tumor cells.(*25, 26*) Given its unique expression, TSHR is recognized as a compelling target for advanced thyroid cancer diagnostics and therapy after thyroidectomy. Overall, there is limited risk of off-tumor adverse effects of targeting TSHR. TSHR has been targeted using biologics including monoclonal antibodies for the treatment of Graves’ disease and thyroid cancer.(*27*) In this study, we generated CART cells targeting TSHR (TSHR-CART) and showed potent, antigen-specific antitumor efficacy against TSHR-expressing cell lines *in vitro* and *in vivo,* as well as in patient-derived xenograft (PDX) mouse models of ATC.

Further, we identified mitogen-activated protein kinase (MAPK) inhibition as a strategy to upregulate TSHR in dedifferentiated thyroid cancers and increase the therapeutic index of TSHR-CART cell therapy. Downregulation of TSHR expression in dedifferentiated thyroid cancer is a potential mechanism of resistance to TSHR-CART cell therapy. MAPK inhibition is an FDA-approved treatment against thyroid cancer and is documented to upregulate TSHR expression in patients with advanced metastatic thyroid cancer.(*28, 29*) Clinical trials for metastatic thyroid cancers have demonstrated that MAPK inhibition induces re-expression of genes necessary for iodine uptake and retention, including TSHR and sodium iodide symporter (NIS).(*30, 31*) Additionally, many thyroid cancers have gene mutations involving the constitutive activation of BRAF, which stimulates MAPK signaling.(*32–34*) Therefore, when BRAF activation is switched off genetically, or its downstream signaling is inhibited with MEK inhibitors (MEKi) or BRAF inhibitors (BRAFi), tumors re-express TSHR and NIS and reaccumulate iodine. This strategy has been employed in the clinic to enhance the response to radioactive iodine therapy.(*29*)

We therefore sought to address antigen escape in ATC by testing the combination of MEK/BRAF small molecule inhibitors with TSHR-CART cell therapy. We found that MEK/BRAF inhibition led to restoration of TSHR expression on thyroid cancer cells and enhancement of CART cell expansion and antitumor activity *in vivo*. This strategy is highly translatable to patients with advanced thyroid cancers who have extremely limited treatment options to date.

## RESULTS

### TSHR as a target for CART cell therapy

To validate TSHR as a target candidate for CART cell therapy in thyroid cancer, we verified its expression on thyroid tissue and lack of expression on other normal tissues by immunohistochemistry (IHC) analysis of specimens from the Mayo Clinic Thyroid Cancer Biobank and on publicly available databases.(*35*) IHC analysis showed that, while TSHR expression was high in normal thyroid tissue as well as multiple thyroid cancer histotypes, ATC samples demonstrated attenuated TSHR expression (**Fig. 1A, B**). We then assessed TSHR expression in a healthy tissue microarray and confirmed that TSHR expression was high in the thyroid and low or absent in other tissues including adult thymus and testis (**Fig. 1C, Supplementary Fig. S1**). Having validated TSHR as a potential antigen to target with CART cell therapy, we designed and generated TSHR-CART cells using a single chain variable fragment isolated from the TSHR autoantibody clone K1-70 and cloned into a lentiviral CAR construct containing 4-1BB and CD3ζ (**Supplementary Fig. S2A**). Healthy donor T cells transduced with CAR-encoded lentiviral particles (**Supplementary Fig. S2B**) demonstrated high CAR expression as assessed by flow cytometry (**Fig. 1D**). TSHR-CART cells displayed comparable CD4:CD8 ratio and phenotype compared to control untransduced T cells (UTD) (**Supplementary Fig. S2C-D**).

**Figure 1.**
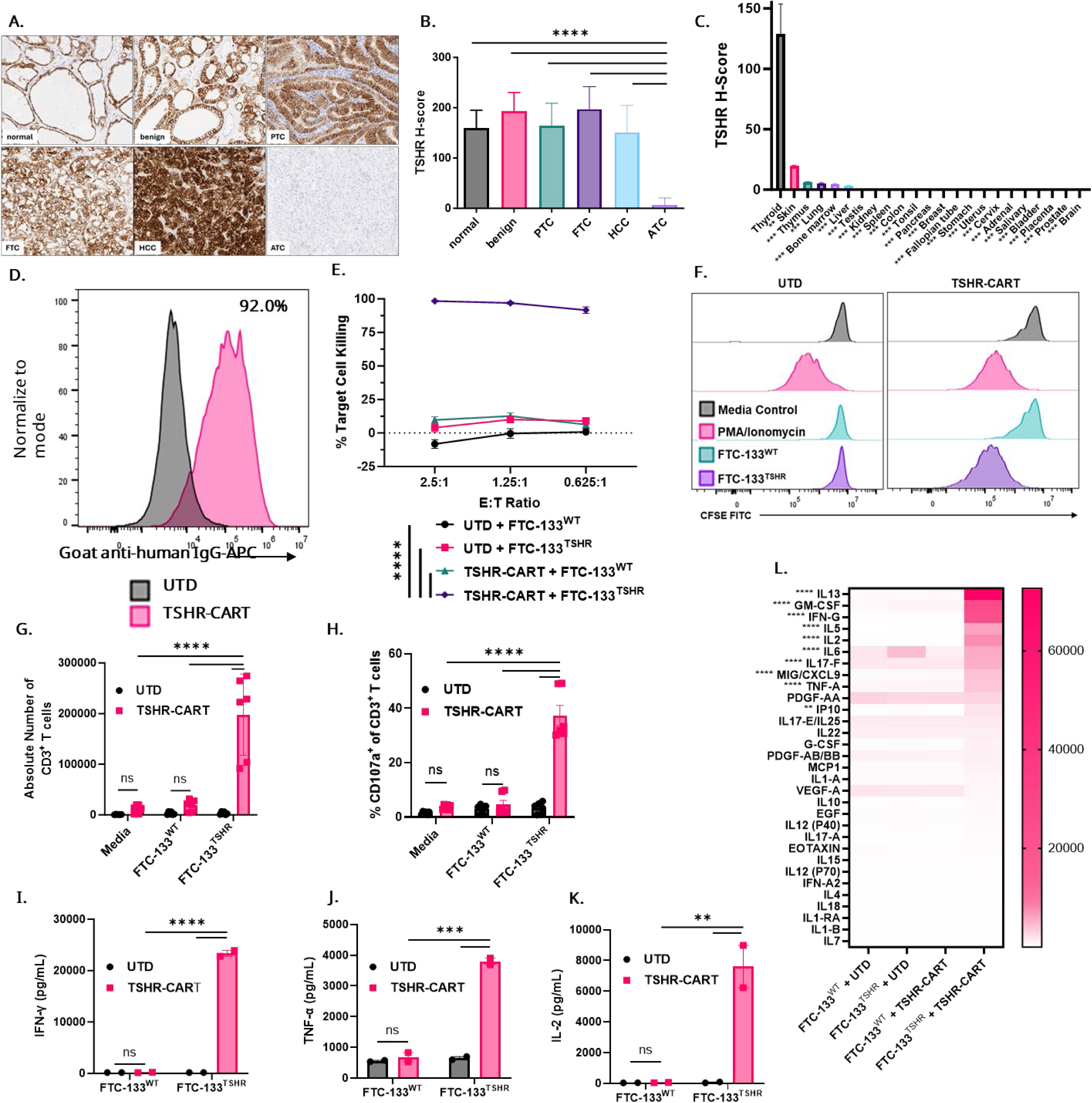
TSHR is a rational target for CART cell therapy, and TSHR-CART cells demonstrate antigen-specific effector functions *in vitro*. **A)** Representative IHC staining shows TSHR expression across normal, benign, and malignant thyroid tissues. PTC: papillary thyroid cancer; FTC: follicular thyroid cancer; HCC: Hurthle cell carcinoma; ATC: anaplastic thyroid cancer. **B)** H-score quantitation of TSHR expression is shown. H-scores provided are based on 0 – 3 IHC scoring and percentage of total areas each score (0x%0 + 1x%1 + 2x%2 + 3x%3 = H Score; 0 – 300 range with statistical analysis (mean and SEM; ****p < 0.0001, one-way ANOVA; normal n = 11, benign n = 9, PTC n = 11, FTC n = 9, HCC n = 37, ATC n = 10). **C)** H-score quantification of normal human tissue stained with TSHR (mean and SEM; **p < 0.01, ***p < 0.001, one-way ANOVA; thyroid n = 7, all other tissues n = 2). **D)** Representative histogram shows CAR expression on TSHR-CART cells as determined by flow cytometry. UTD T cells were used as a negative control. **E)** Luciferase^+^ TSHR^+^ or TSHR^-^FTC-133 target cells were co-cultured with either UTD or TSHR-CART cells at varying effector-to-target ratios, and cytotoxicity was measured via bioluminescence after 48 hours (mean and SEM; ***p < 0.001, ****p < 0.0001, two-way ANOVA; 3 biological replicates). **F-G)** CFSE-labeled UTD or TSHR-CART cells were co-cocultured with media alone (negative control), PMA/ionomycin (positive control), or TSHR^+^ or TSHR^-^ FTC-133 target cells. After five days, T cell proliferation was assessed by flow cytometry using CFSE staining (F) or absolute counts of CD3^+^ cells (G) (mean and SEM; ****p < 0.0001, two-way ANOVA; 3 biological replicates). **H)** UTD or TSHR-CART cells were-cocultured with TSHR^+^ FTC-133 target cells at a 1:5 ratio for four hours, and degranulation was then assessed by flow cytometry of CD3^+^ CD107a^+^ cells (mean and SEM; ****p < 0.0001, two-way ANOVA; 2 biological replicates). **I-K)** UTD or TSHR-CART cells were-cocultured with TSHR^-^ or TSHR^+^ FTC-133 target cells at a 1:1 ratio for three days, and supernatants were analyzed for human cytokines via a 38-multiplex assay (mean and SEM; **p < 0.01, ***p < 0.001, ****p < 0.0001, two-way ANOVA; 2 biological replicates). **L)** Heat map of additional analytes from the 38-plex assay (median; **p < 0.01, ****p < 0.0001, two-way ANOVA; 2 biological replicates).

### TSHR-CART cells exhibit potent and specific antitumor activity *in vitro*

We next examined the antigen-specific activation of TSHR-CART cells *in vitro* and *in vivo*. TSHR-CART cells or UTD were incubated with the follicular thyroid cancer cell line, FTC-133, which was transduced to overexpress TSHR; TSHR was transduced in these cells as thyroid cancer cell lines and patient samples downregulate TSHR expression after extended monolayer cell culture.(*36–39*) While TSHR-CART cells were not activated against wildtype (TSHR^-^) FTC-133 cells, TSHR-CART cells demonstrated profound antigen-specific effector functions against TSHR^+^ target cells *in vitro*, including cytotoxicity against target cells (**Fig. 1E**) as well as TSHR-CART proliferation (**Fig. 1F-G, Supplementary Fig. S3A**), degranulation (**Fig. 1H**), and cytokine production (**Fig. 1I-L**). Similar results were found for wildtype (TSHR^-^) and TSHR^+^ THJ-529, a PDTC cell line (**Supplementary Fig. S3B-H**).

### TSHR-CART cells show durable antitumor activity in xenograft HCC models

We then assessed TSHR-CART cell activity in a xenograft model of HCC. We first measured TSHR expression via IHC on biopsies obtained from patients with normal thyroid tissue, primary HCC with minimal or extensive invasion, locally recurrent HCC, and metastatic HCC. Strong TSHR expression was maintained in recurrent and metastatic lesions as well as primary tumors (**Fig. 2A-B**). Having verified that HCC retains TSHR expression in patient samples and thus providing a strong rationale for the use of TSHR-CART cell monotherapy, we then assessed the *in vivo* activity of TSHR-CART cells against the TSHR^+^ HCC cell line, XCT.UC1 as a proof of concept. Non-obese diabetic severe combined immunodeficiency IL2rγ^-^/^-^ (NSG) mice were subcutaneously engrafted with 1 x 10^7^ XTC.UC1, which retains high endogenous TSHR expression *in vivo*, and treated with 1 x10^7^ UTD or TSHR-CART cells when tumors reached ∼100mm^3^. Mice in remission were rechallenged with 1 x 10^7^ XTC.UC1 cells subcutaneously in the opposite flank (**Fig. 2C**). Serial caliper measurements revealed significant antitumor efficacy of TSHR-CART cells compared to UTD control; by Day 50 post-engraftment, all UTD-treated mice had reached humane endpoints, while all TSHR-CART-treated mice were in remission or had small tumors (**Fig. 2D-E**). TSHR-CART-treated mice in remission were then rechallenged on Day 51 with XTC.UC1 in the opposite flank and were followed for tumor growth along with a new group of untreated XTC.UC1-bearing mice as controls. Again, TSHR-CART cells displayed potent antitumor activity; both original and rechallenged tumors were significantly suppressed compared to untreated controls (**Fig. 2F**). Peripheral blood sampling at 21 days post-tumor rechallenge revealed significantly higher numbers of human CD3^+^ T cells in TSHR-CART-treated mice compared to controls (**Fig. 2G**). These data demonstrated that TSHR-CART cells exerted durable tumor control in HCC, an aggressive disease setting with no current standard of care.

**Figure 2.**
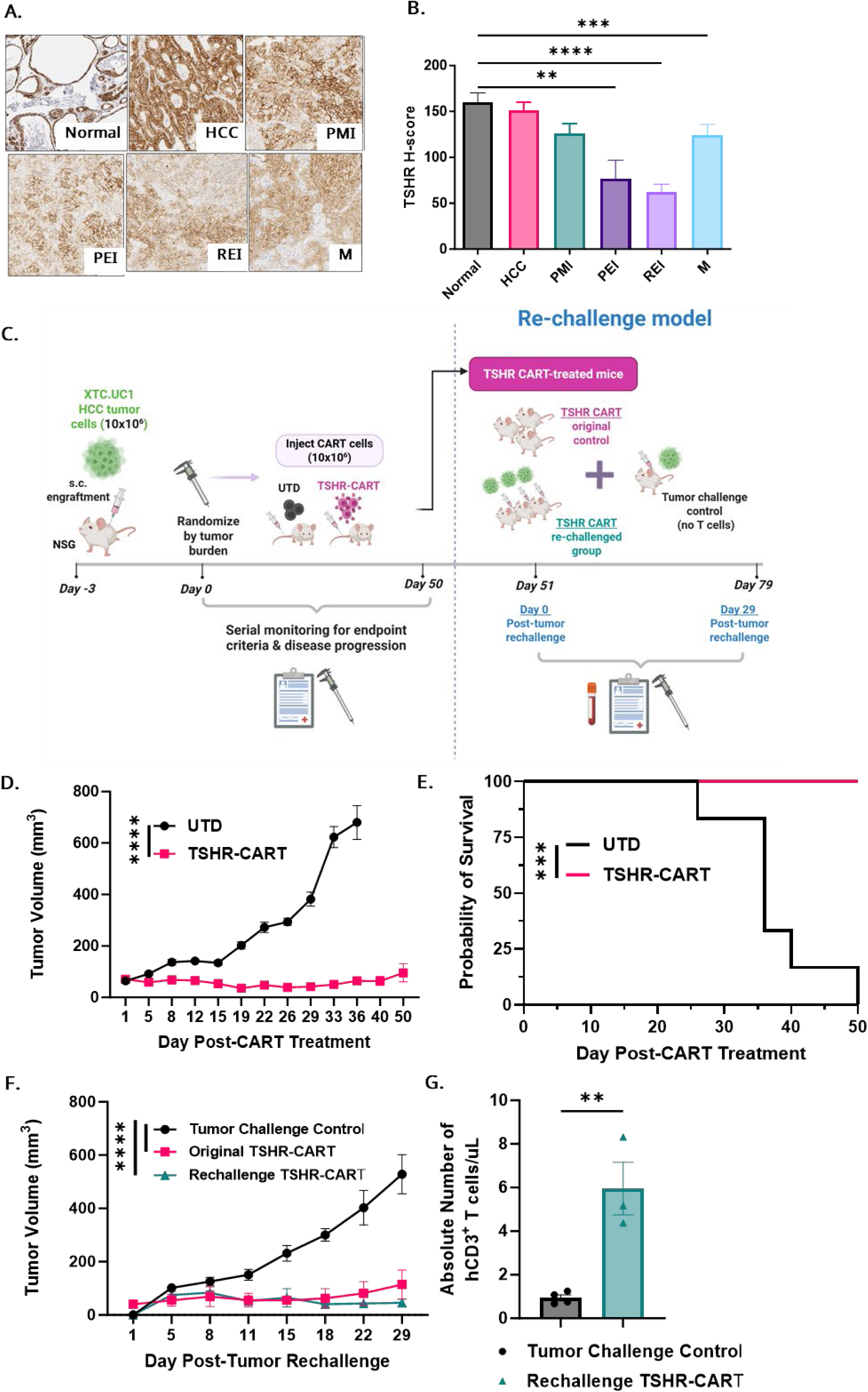
TSHR-CART cells exhibit potent antitumor activity in vivo against Hurthle cell carcinoma. **A)** Representative IHC staining of TSHR expression across normal and HCC tissues. Hurthle cell carcinoma: HCC; primary tumor with minimal invasion: PMI; primary tumor with extensive invasion: PEI; recurrent tumor with extensive invasion: REI; metastatic tumors: M. **B)** TSHR expression was quantified using H-Score (based on 0 – 300 IHC scoring and percentage of total areas each score, 0 – 300 range with statistical analysis) (mean and SEM; **p < 0.01, ***p < 0.001, ****p <0.0001, one-way ANOVA. Normal n=11, HCC n=37, PMI n=26, PEI n=11, REI n=34, M n=16). **C)** Experimental schema of a HCC xenograft model. NSG mice were subcutaneously engrafted with TSHR^+^ XTC.UC1 thyroid cancer cells in Matrigel. Three days later, after engraftment was confirmed, mice were randomized according to tumor volume to receive either UTD (10 X 10^6^) or TSHR-CART cells (10 x 10^6^) intravenously. Mice were monitored for well-being, tumor volume, and overall survival. At day 51, mice treated with TSHR-CART cells which achieved complete remission were rechallenged subcutaneously with XCT.UC1 cells in the opposite flank. Additionally, an untreated group was engrafted with XCT.UC1 cells and included as a negative control. Mice were monitored for well-being, tumor volume, T cell expansion, and overall survival. **D)** Tumor volume was assessed via serial caliper measurements (mean and SEM; ****p < 0.0001, two-way ANOVA; n=6 mice per group). **E)** Kaplan-Meier survival curve (***p < 0.001, Log-rank test; n=6 mice per group). **F)** Tumor volume as assessed by serial caliper measurements (mean and SEM; ****p < 0.0001, two-way ANOVA; n=6 mice per group). **G)** Peripheral blood was collected 21 days after tumor rechallenge and assessed by flow cytometry using absolute counts of hCD3^+^ cells (mean and SEM; **p < 0.01, two-tailed unpaired t-test; n = 3 mice per group).

### TSHR-CART cells demonstrate strong antitumor effects in TSHR^+^ PDX models of PDTC

Next, we evaluated the antitumor activity of TSHR-CART cells in a dose escalation model in NSG mice. Here, we utilized aggressive, BRAF-mutant, patient-derived PDTC cells, THJ-529T, which were transduced to overexpress TSHR. Mice were subcutaneously injected with 2.5 x 10^6^ THJ-529T cells, and engraftment was confirmed by caliper measurement of tumor masses. When tumors reached ∼100mm^3^, mice were randomized to treatment with 5 x 10^6^ UTD, 5 x 10^6^ TSHR-CART cells, or 1 x 10^7^ TSHR-CART cells (**Fig. 3A**). Treatment with TSHR-CART cells led to significant and dose-dependent tumor control (**Fig. 3B-E, Supplemental Fig. S4A-C**), improved survival outcomes (**Fig. 3F, Supplemental Fig. S4D**), increased tumor-infiltrated human T cells (**Fig. 3G-H**), and increased human T cell proliferation in the peripheral blood (**Supplemental Fig. S4E**). Having shown that the 1 x 10^7^ TSHR-CART dose yielded superior antitumor activity compared to the 5 x 10^6^ TSHR-CART dose, we did not further pursue testing of the 5 x 10^6^ TSHR-CART dose.

**Figure 3.**
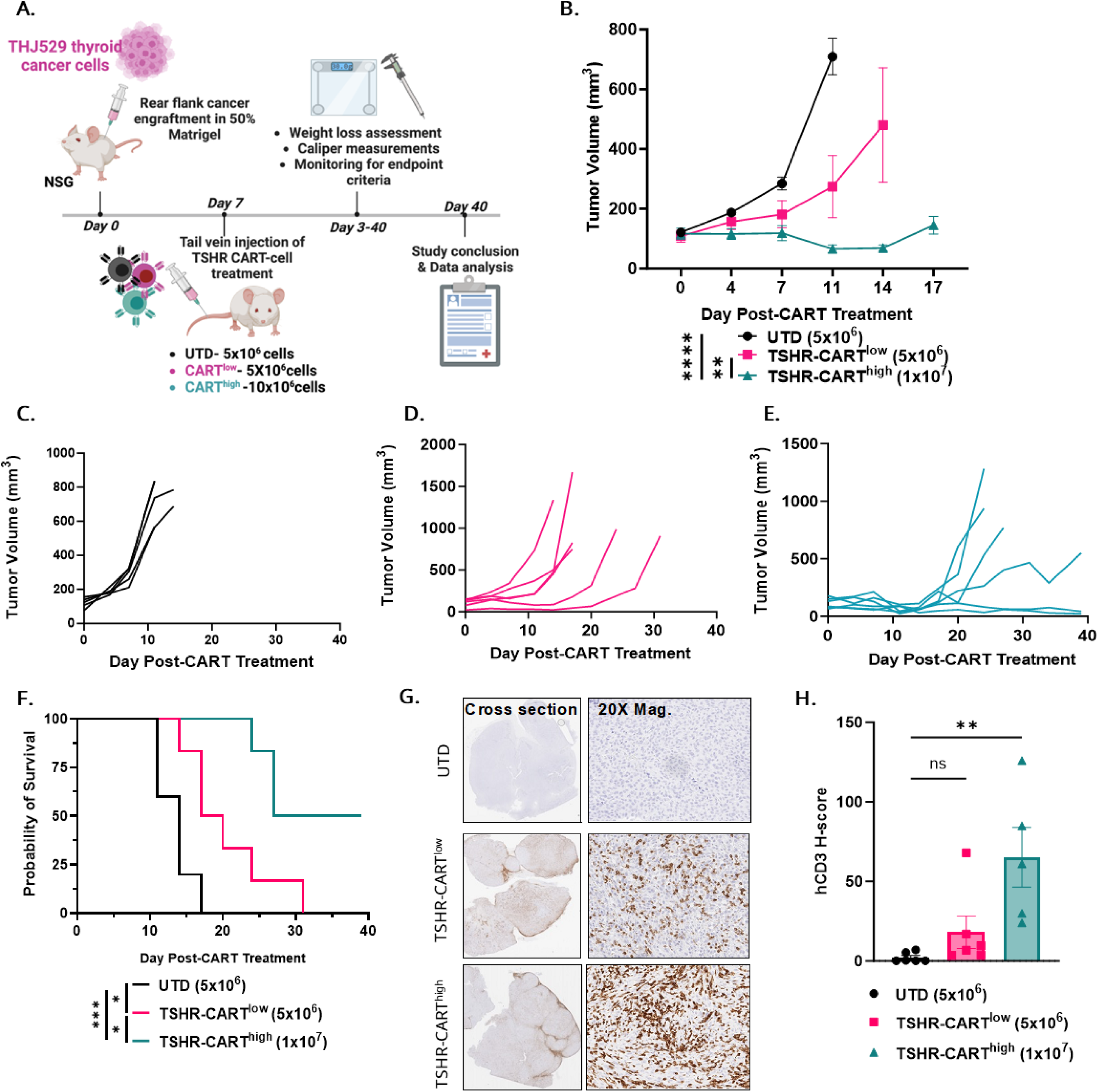
TSHR-CART cells exhibit dose-dependent antitumor activity in PDX models of TSHR^+^ PDTC. **A)** Experimental schema of a PDTC xenograft model. NSG mice were subcutaneously engrafted with TSHR^+^ THJ-529 thyroid cancer cells in Matrigel. One week later, after engraftment was confirmed, mice were randomized according to tumor volume and received either UTD (5 x 10^6^) or TSHR-CART cells (5 x 10^6^ or 1 x 10^7^) intravenously. Mice were monitored for well-being, tumor volume, and overall survival. **B)** Tumor volume was assessed via serial caliper measurements (mean and SEM; **p < 0.01, ****p < 0.0001, two-way ANOVA; n=5-6 mice per group). **C-E)** Individual tumor growth curves are shown for each treatment group, including UTD (**C**), TSHR-CART^low^ (**D**), and TSHR-CART^high^ (**E**). N = 5-6 mice/group. **F)** Kaplan-Meier survival curve (*p < 0.05, ***p < 0.001, Log-rank test; n=5-6 mice per group). **G)** Representative IHC staining shows human CD3 expression in thyroid tumors excised from mice in each treatment group. **H)** H-score quantitation of human CD3 expression is shown. H-scores provided are based on 0 – 3 IHC scoring and percentage of total areas each score (0x%0 + 1x%1 + 2x%2 + 3x%3 = H Score; 0 – 300 range with statistical analysis) (mean and SEM; **p < 0.01, one-way ANOVA; n = 5-6 mice per group).

### TSHR is downregulated in ATC but is restored with MAPK inhibition

ATC cells have been associated with de-differentiation and downregulation of TSHR expression, and we showed that ATC significantly downregulated TSHR expression compared to normal thyroid tissues or more differentiated thyroid cancers (**Fig. 1A, B**). This observed downregulation raises concerns for antigen escape and subsequent resistance to TSHR-CART cell therapy. We first verified that the MEKi, trametinib, restored TSHR expression in an ATC PDX proof-of-concept model. In this model, 5mm^3^ Th-560 ATC PDX tissue was surgically implanted subcutaneously in NSG mice. When tumors reached approximately 100mm^3^, mice were randomized to daily treatment with either vehicle control or MAPK inhibitors at the indicated doses until the experimental endpoint (**Fig. 4A**). The administration of MEKi (trametinib) resulted in significant dose-dependent slowing of tumor growth (**Fig. 4B**) as well as TSHR re-expression which inversely correlated with pERK expression (**Fig. 4C**) after one week of daily trametinib treatment compared to placebo treatment. Because many ATC tumors are BRAF-mutant, we performed a similar experiment in which mice received either vehicle control or the dual MEKi/BRAFi R05126766 at the indicated doses. Treatment with R05126766 resulted in a significant dose-dependent slowing of tumor growth (**Fig. 4D**), and pERK expression inversely correlated with TSHR expression (**Fig. 4E**) after one week of daily R05126766 treatment compared to placebo treatment. Similarly, treatment with combination of MEKi (trametinib) and BRAFi (dabrafenib) resulted in significantly increased TSHR expression in ATC PDX models (**Fig. 4F-G**). While modest antitumor effects of MEKi and/or BRAFi were observed, the main purpose of these experiments was to assess TSHR upregulation upon MEKi and/or BRAFi treatment in PDX models of ATC. Collectively, these data support the therapeutic strategy of MEKi and BRAFi treatment to re-express TSHR and thereby reduce ATC resistance to TSHR-CART cell therapy.

**Figure 4.**
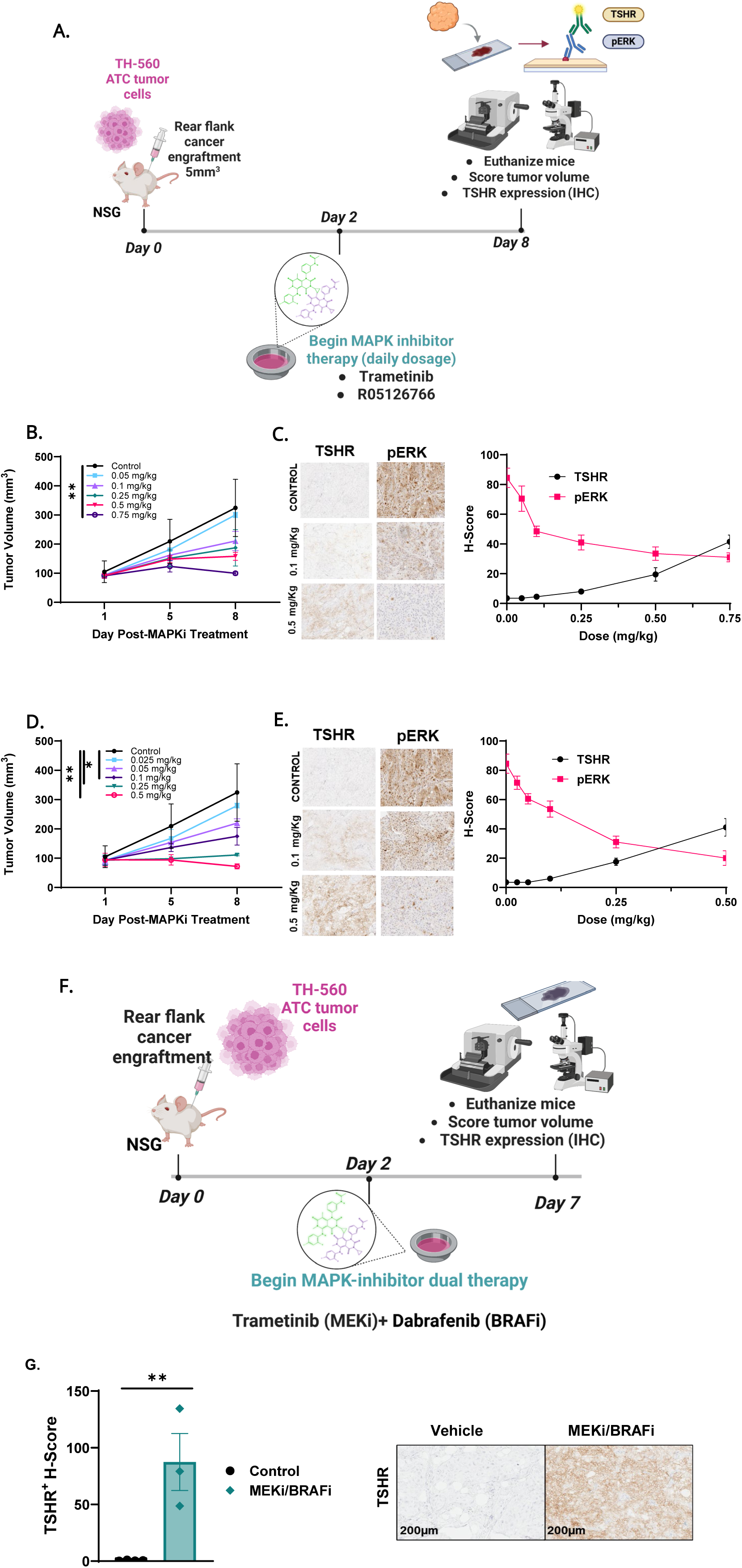
Treatment of ATC PDX tumors with MAPK inhibitors restores TSHR expression. **A)** Experimental schema of an ATC PDX model. NSG mice were subcutaneously engrafted with TH-560 ATC PDX tumors. When tumors reached approximately 100 mm^3^, mice were orally treated with MAPK inhibitors daily and serially monitored for tumor volume. On day 8, mice were sacrificed, and tumors were harvested and assessed for TSHR expression by IHC. **B)** Tumor volumes were assessed via serial caliper measurements in mice treated with trametinib at the indicated doses (mean and SEM; **p < 0.01, two-way ANOVA; n=2 mice per group). **C)** Representative IHC staining (left) shows human TSHR and pERK expression in thyroid tumors excised from mice in each treatment group. H-scores (right) provided are based on 0 – 3 IHC scoring and percentage of total areas each score (0x%0 + 1x%1 + 2x%2 + 3x%3 = H Score; 0 – 300 range with statistical analysis. Mean and SEM; n = 2 mice per group). **D)** Tumor volumes were assessed via serial caliper measurements in mice treated with RO5126766 at the indicated doses (mean and SEM; *p < 0.05, **p < 0.01, two-way ANOVA; n=2 mice per group). **E)** Representative IHC staining (left) and H-score quantitation (right) show human TSHR and pERK expression in thyroid tumors excised from mice in each treatment group. H-scores provided are based on 0 – 3 IHC scoring and percentage of total areas each score (0x%0 + 1x%1 + 2x%2 + 3x%3 = H Score; 0 – 300 range with statistical analysis. Mean and SEM; n = 2 mice per group). **F)** Experimental schema of an ATC PDX model. NSG mice were subcutaneously engrafted with Th-560 ATC PDX tumors. When tumors reached approximately 100 mm^3^, mice were orally treated with MAPK inhibitors daily and serially monitored for tumor volume. On Day 7, mice were sacrificed, and tumors were harvested and assessed for TSHR expression by IHC. **G)** Representative IHC staining (top) and H-score quantitation (bottom) show human TSHR expression in thyroid tumors excised from mice in each treatment group. H-scores provided are based on 0 – 3 IHC scoring and percentage of total areas each score (0x%0 + 1x%1 + 2x%2 + 3x%3 = H Score; 0 – 300 range with statistical analysis. Mean and SEM; **p <0.01, unpaired two-tailed t-test, n = 3-4 mice per group).

### MAPK inhibition upregulates TSHR and sensitizes TSHR-CART cell therapy to dedifferentiated thyroid cancer *in vitro*

We then tested if pretreatment with MEKi or MEKi/BRAFi enhanced TSHR-CART cell activity against dedifferentiated thyroid cancer cell lines *in vitro.* The PDTC cell line, THJ-529, was cocultured with trametinib alone or trametinib plus dabrafenib (10nM or 1µM for trametinib or dabrafenib). After four days, THJ-529 cells were washed and assessed for TSHR expression by flow cytometry. THJ-529 cells demonstrated modest but consistently increased TSHR expression after treatment with MEKi or MEKi/BRAFi compared to vehicle-treated cells (**Fig. 5A**). THJ-529 cells were then washed to remove residual MAPK inhibitor and plated with TSHR-CART cells for five additional days, at which point TSHR-CART cell proliferation was assessed via flow cytometry. TSHR-CART cells demonstrated significantly more proliferation in response to THJ-529 cells which were pretreated with trametinib or trametinib plus dabrafenib compared to vehicle control-treated THJ-529 cells (**Fig. 5B**). Treatment with higher or more prolonged MAPK inhibitor doses to further increase TSHR expression resulted in tumor cell death to the point where post-coculture analyses could not be conducted; however, the modest TSHR upregulation observed with low doses of MAPK inhibitors resulted in significantly increased TSHR-CART response. We then assessed the sensitivity of TSHR-CART cells to low expression levels of TSHR. We generated a series of TSHR^+^ FTC-133 cells with varied levels of TSHR expression. TSHR^+^ FTC-133 cells had TSHR levels ranging from ∼40,000 to 300,000 TSHR per cell, compared to ∼6,000 TSHR per cell for our ATC PDX model, THJ560, and ∼1,000 TSHR per cell for wildtype FTC-133 (**Fig. 5C-D**). TSHR-CART cells demonstrated potent cytotoxicity against all TSHR^+^ FTC-133 cell lines, with only the wildtype FTC-133 cells showing resistance to TSHR-CART cell killing (**Fig. 5E**). These findings indicated that low to moderate TSHR upregulation via MAPK inhibition leads to robust antitumor activity of TSHR-CART cells.

**Figure 5.**
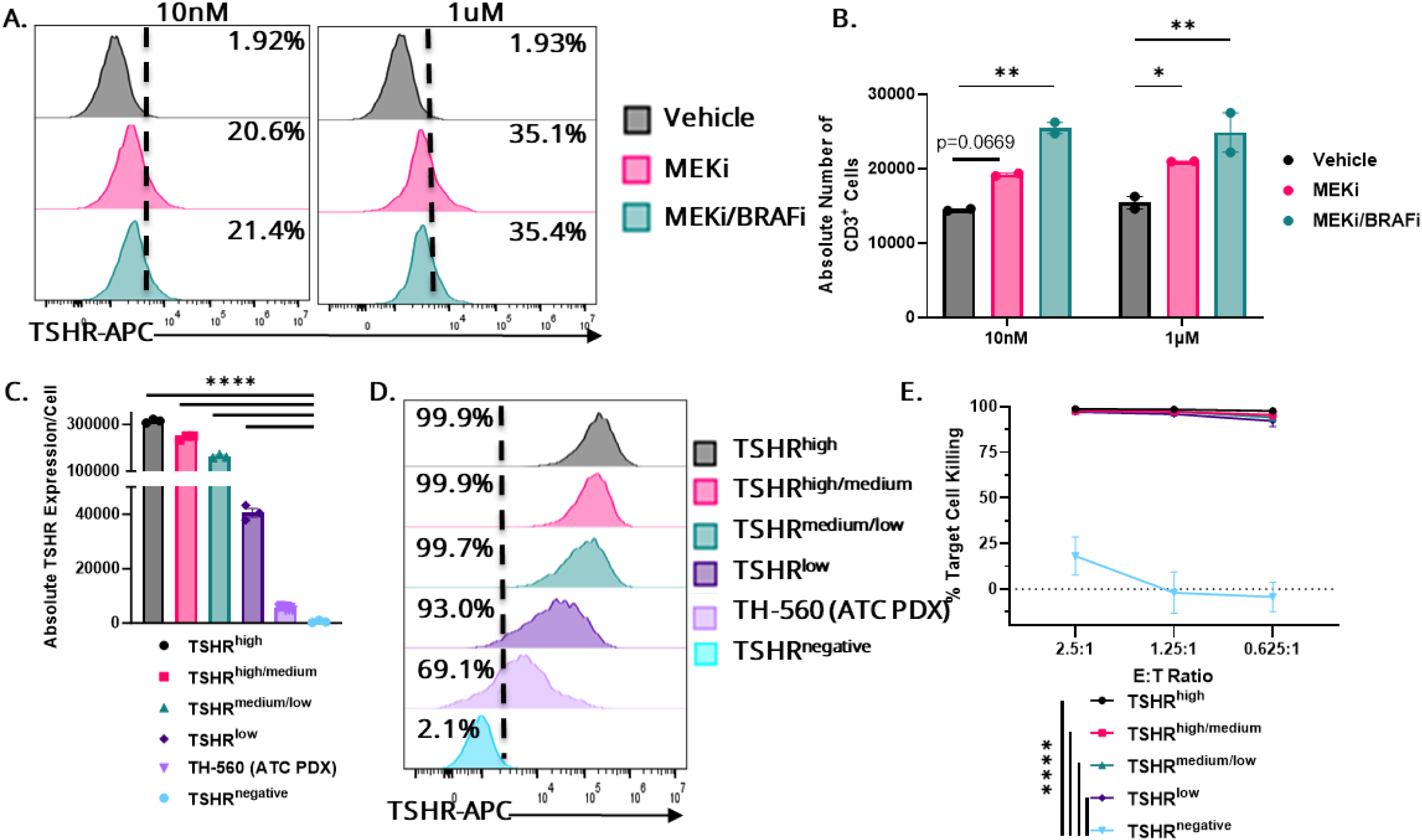
TSHR-CART cells are sensitive to moderate upregulation of TSHR on target cells. **A)** THJ-529 cells were cocultured with 10 nM or 1µM vehicle control, trametinib, or both trametinib and dabrafenib. After four days, THJ-529 cells were trypsinized and flowed for TSHR expression. Representative histogram shown. **B)** After four days of MAPK inhibitor co-culture, THJ-529 cells were trypsinized and plated with TSHR-CART cells at a 1:1 ratio and assessed for proliferation five days later as determined by flow cytometry of absolute number of CD3^+^ cells (mean and SEM; *p < 0.05, **p < 0.01, two-way ANOVA; 1 biological replicate). **C)** Assessment of absolute TSHR expression levels in wildtype or lentivirally transduced TSHR^+^ FTC-133 tumor cells and wildtype PDX tumors, as determined by flow cytometry using absolute counting beads (mean and SEM; ****p < 0.0001, one-way ANOVA; n= 3-9 technical replicates). **D)** Representative histogram shows different levels of TSHR expression on tumor cells as determined by flow cytometry. **E)** Percentage of killing measured via bioluminescent imaging after 48 hours of co-culture at different E:T ratios of TSHR-CART cells and luciferase^+^ FTC-133 tumor cells expressing various levels of TSHR (mean and SEM; ****p < 0.0001, two-way ANOVA; 2 biological replicates).

### MAPK inhibition does not negatively impact TSHR-CART cell activity

Therefore, we aimed to evaluate MEK and BRAF inhibition as a strategy to upregulate TSHR expression on thyroid cancer cells and increase the therapeutic index of TSHR-CART cell therapy. Given that T cells are shown to utilize MAPK signaling, we first tested for potential detrimental effects of MEKi or MEKi/BRAFi on CART cell activity. TSHR-CART proliferation (**Fig. 6A**), killing (**Fig. 6B**), degranulation (**Supplemental Fig. S5A**), and intracellular cytokine production (**Supplemental Fig. S5B**) were not affected by the addition of trametinib alone or trametinib plus dabrafenib compared to media control. We next assessed the effects of prolonged MAPK inhibition on TSHR-CART function through an exhaustion assay developed with repeated CART activation.(*40*) Here, TSHR-CART cells were repeatedly stimulated with irradiated TSHR^+^ FTC-133 cells along with vehicle control, 10nM trametinib, or 10nM trametinib plus 10nM dabrafenib. After one week of chronic stimulation, TSHR-CART cells were isolated and cocultured with fresh TSHR^+^ FTC-133 cells and vehicle control, 10nM trametinib, or 10nM trametinib plus dabrafenib for proliferation and cytotoxicity assays (**Supplemental Fig. S5C**). Here as well, the addition of MAPK inhibitors did not appear to negatively impact TSHR-CART cell proliferation (**Fig. 6C, Supplemental Fig. S5D**) or killing (**Fig. 6D**).

**Figure 6.**
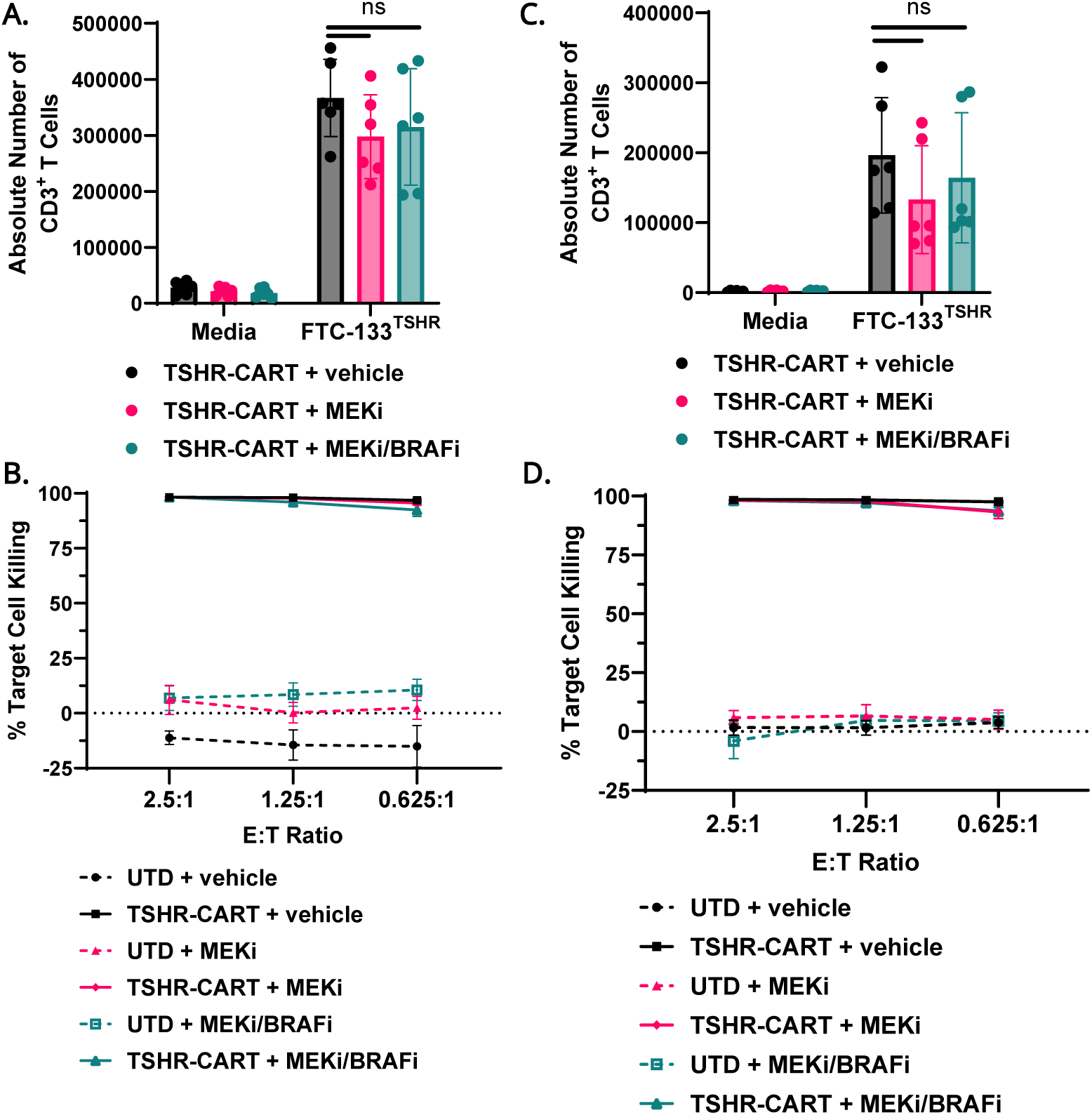
TSHR-CART effector functions remain intact after the addition of MAPK inhibitors. **A**) TSHR-CART cells were co-cocultured with either media alone (negative control) or TSHR^+^ FTC-133 target cells at a 1:1 ratio, along with vehicle control, 10nM trametinib, or 10nM both trametinib and dabrafenib. After five days, T cell proliferation was assessed by flow cytometry using absolute counts of CD3^+^ cells (mean and SEM; two-way ANOVA; 2 biological replicates). **B)** TSHR-CART cells were-cocultured with luciferase^+^ TSHR^+^ FTC-133 target cells at various E:T ratios, along with vehicle control, 10nM trametinib, or 10nM both trametinib and dabrafenib. After 48 hours, target cell killing was assessed by bioluminescence (mean and SEM; two-way ANOVA; 2 biological replicates). **C-D)** In chronic stimulation assays, TSHR-CART cells were co-cocultured with either media alone (negative control) or irradiated TSHR^+^ FTC-133 target cells at a 1:1 ratio, along with vehicle control, 10nM trametinib, or 10nM both trametinib and dabrafenib. Irradiated target cells and MAPK inhibitor-containing media were replenished every other day for one week. Then, TSHR-CART cells were isolated with CD4 and CD8 beads and plated in proliferation (**C**) or cytotoxicity (**D**) assays as described above (mean and SEM; two-way ANOVA; 2 biological replicates).

### Combination of TSHR-CART cells plus MAPK inhibitors yields superior TSHR-CART cell activity

Having shown that the addition of MAPK inhibitors was not detrimental to CART function, we next studied the kinetics of TSHR expression after MEKi or MEKi/BRAFi to determine the longevity of TSHR re-expression after treatment with MAPK inhibitors. Mice were implanted with 5mm^3^ Th-560 ATC PDX tumors. When tumors reached ∼100mm^3^, mice were treated daily with either vehicle control or with 1.5 mg/kg dual MEKi/BRAFi (R05126766) plus 1 mg/kg BRAFi (dabrafenib). TSHR expression was assessed by IHC and was found to be significantly elevated after 7 days of MAPK inhibitor treatment compared to vehicle control. However, TSHR expression was subsequently lost within two days after stopping MAPK inhibitor treatment (**Supplemental Fig. S6A-B**).

Given these results, we compared different dosing regimens of MEKi/BRAFi in combination with TSHR-CART cell therapy in ATC PDX models to determine if sequential or concurrent MAPK inhibitor administration yielded superior CART antitumor activity. Mice were subcutaneously implanted with 5mm^3^ Th-560 ATC PDX tumors. When tumors reached ∼100mm^3^, mice were randomized to daily treatment with vehicle control, 0.25 mg/kg trametinib (concurrently administered until experimental endpoints or stopped after 7 days), or 0.25 mg/kg trametinib plus 2 mg/kg dabrafenib (concurrently administered until experimental endpoints or stopped after 7 days). Mice treated with vehicle control were engrafted with tumor one week later than mice treated with MAPK inhibitors to ensure equal tumor sizes across all groups when UTD or TSHR-CART cells were administered. As a further dose escalation, mice were then given 2 x 10^7^ UTD or TSHR-CART cells via tail vein injection. Tumor volume, body weight, and body condition were inspected to monitor disease progression and to assess for endpoint criteria (**Fig. 7A**). Mice treated with concurrent MEKi or concurrent MEKi plus BRAFi in combination with TSHR-CART cells (CART + MEKi and CART + MEKi/BRAFi, respectively) showed superior antitumor efficacy (**Fig. 7B-C, Supplementary Fig. S7A-F, Supplementary Fig. S8A-F**). We then performed a followup experiment using 1 x 10^7^ UTD or TSHR-CART cells. Here, mice with concurrent MAPK inhibitor administration plus TSHR-CART cells demonstrated durable antitumor control lasting beyond two months (**Supplementary Fig. S8G**). Given these results, we used the 1 x 10^7^ TSHR-CART dose in subsequent experiments. These findings indicate that administering MAPK inhibitors concurrently with TSHR-CART cells enhances the antitumor activity of TSHR-CART cell therapy.

**Figure 7.**
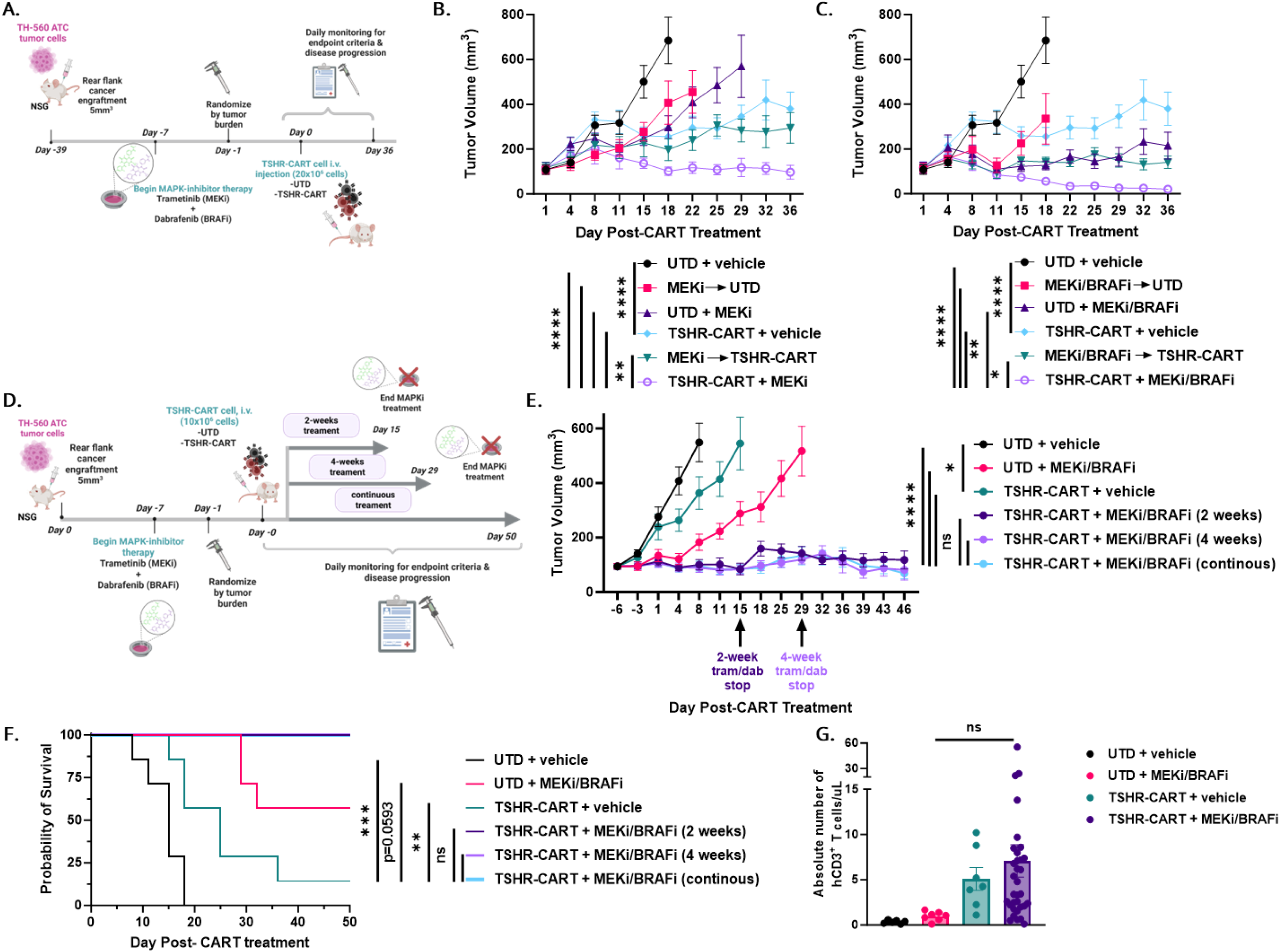
TSHR-CART cells show the most potent antitumor activity when combined with MAPK inhibitor administration. **A)** Experimental schema of an ATC PDX model. NSG mice were subcutaneously engrafted with Th-560 ATC PDX tumors. When tumors reached approximately 100 mm^3^, mice were orally treated with MAPK inhibitors (0.25 mg/kg trametinib alone or with 2 mg/kg dabrafenib) daily for either seven days prior to TSHR-CART administration or concurrently (until endpoints were reached) and serially monitored for tumor volume. Mice were then randomized by tumor volume and treated with either UTD or TSHR-CART cells (20x10^6^ i.v.). Mice were monitored for well-being and tumor volume. **B)** Tumor volumes were assessed via serial caliper measurements in mice treated with UTD or TSHR-CART with vehicle control, with 7 days of trametinib, or with concurrent trametinib at the indicated doses (mean and SEM; **p < 0.01, ****p < 0.0001, two-way ANOVA; n=7 mice per group). **C)** Tumor volumes were assessed via serial caliper measurements in mice treated with UTD or TSHR-CART with vehicle control, with 7 days of trametinib plus dabrafenib, or with concurrent trametinib plus dabrafenib at the indicated doses (mean and SEM; *p < 0.05, **p < 0.01, ****p < 0.0001, two-way ANOVA; n=7 mice per group). **D)** Experimental schema of an ATC PDX model. NSG mice were subcutaneously engrafted with Th-560 ATC PDX tumors. When tumors reached approximately 100 mm^3^, mice were orally treated with MAPK inhibitors (0.25 mg/kg trametinib plus 2 mg/kg dabrafenib) daily for seven days monitored for tumor volume. After 7 days of MAPK inhibitor treatment, mice were randomized by tumor volume to be treated with either UTD or TSHR-CART cells (10x10^6^ i.v.). After UTD or TSHR-CART injection, mice were separated to vehicle control or MAPK inhibitors (for 2 weeks, for 4 weeks, or continuously until experimental endpoints were reached. **E)** Mice treated with TSHR-CART cells in combination with MAPK inhibitors showed superior tumor volume control regardless of the duration of MAPK inhibitor treatment. Tumor volumes were assessed via caliper measurements in mice treated with UTD or TSHR-CART cells in combination with vehicle control or MAPK inhibitors (mean and SEM; ns = not significant, *p < 0.05, ****p < 0.0001, two-way ANOVA; n=7 mice per group. **F)** Kaplan-Meier survival curve (**p < 0.01, ***p < 0.001, Log-rank test; n=7 mice per group). **G)** *In vivo* TSHR-CART cell expansion was measured by absolute count of hCD45^+^ hCD3^+^ cells per µL of blood via flow cytometry analysis (mean and SEM; ns = not significant, one-way ANOVA; n = 7-33 mice per group).

We next assessed how long MAPK inhibitors needed to be administered concurrently with TSHR-CART cell therapy for enhanced antitumor efficacy. Here, mice treated with UTD or TSHR-CART cells were given vehicle control, MEKi/BRAFi for two weeks after CART administration, MEKi/BRAFi, for four weeks after CART administration, or continuous MEKi/BRAFi after CART administration until experimental endpoints were reached (**Fig. 7D**). Mice receiving the combination of TSHR-CART and MAPK inhibitors showed superior tumor control and survival, regardless of the duration of concurrent MAPK inhibitors (**Fig. 7E-F, Supplementary Fig. S9A-F**). This indicates that using MAPK inhibitors in combination with TSHR CART cells for 2 weeks is sufficient to induce TSHR upregulation and potent TSHR activity. Peripheral blood analysis at day 14 post-CART injection showed comparable human T cell expansion in mice treated with TSHR-CART cells plus vehicle control compared to mice treated with TSHR-CART cells plus MAPK inhibitors (**Fig. 7G**).

## DISCUSSION

Here, we describe the development and evaluation of a highly translational CART cell therapy for the treatment of poorly differentiated, anaplastic, and metastatic thyroid cancers. TSHR is a rational target for CART cell therapy, given that it is widely expressed in thyroid cancers and rarely expressed outside the thyroid. We subsequently generated and tested TSHR-CART cells, which demonstrated antigen-specific activity against TSHR-expressing target cells *in vitro* and *in vivo*. HCC represents ∼4% of thyroid cancers and is typically radioiodine-resistant for metastatic disease, retains TSHR expression, and currently has no FDA-approved therapy.(*7*) Our data demonstrate that TSHR-CART cell therapy resulted in durable remissions *in vivo* and may represent a treatment option for this subset of patients.

De-differentiated and anaplastic thyroid cancers can exhibit significant downregulation of thyroid developmental genes, including TSHR.(*41, 42*) This phenomenon poses a formidable hurdle to effective TSHR-CART treatment. Increasing evidence indicates that TSHR downregulation and de-differentiation is mediated by MAPK signaling.(*28, 29, 34, 43, 44*) The majority of thyroid cancers have activating oncogenic gene mutations involving PI3K, RET, RAS, and BRAF.(*32*) Constitutive activation of these proteins stimulates MAPK signaling, inhibiting the expression of thyroid hormone biosynthesis genes.(*33, 34*) Therefore, when BRAF activation is switched off genetically or its downstream signaling is inhibited with MEKi or BRAFi, tumors re-express TSHR and reaccumulate iodine. Such a strategy has been employed in the clinic to enhance the response to radioactive iodine therapy.(*30*) Recent findings from patients with thyroid cancer who have been treated with MEKi indicate a dramatic upregulation of TSHR within both primary and secondary thyroid cancer tumors.(*30, 45*) Additionally, MEKi, alone or in combination with BRAFi, demonstrate significant antitumor activity in metastatic thyroid cancer and are FDA-approved for ATC.(*46*) We therefore tested the combination of TSHR-CART cell therapy plus MAPK inhibition to upregulate TSHR expression in the tumor and enhance CART efficacy, demonstrating restoration of TSHR expression and improved CART antitumor effects in mice receiving combination therapy compared to mice receiving CART monotherapy.

Major limitations to effective CART cell therapy in solid tumor settings include appropriate target antigen selection, antigen-negative escape, and lack of T cell infiltration into the tumor site. The treatment strategy we have outlined here has strong potential to overcome these limitations, as TSHR is nearly uniquely and universally expressed in thyroid tumors, MAPK inhibition induces TSHR re-expression, and TSHR-expressing advanced thyroid tumors are commonly infiltrated by immune cells, including T cells.(*9*)

Regarding the potential for on-target off-tumor toxicity, consistent with publicly available datasets, our tissue microarray demonstrates that TSHR expression is mainly limited to the thyroid gland. Furthermore, our CAR is cross reactive with mouse TSHR and we did not observe any toxicities in our mouse experiments. As patients with aggressive thyroid cancers will have undergone previous thyroidectomy, there will be no healthy thyroid tissue remaining upon treatment with TSHR-CART cells. There have also been reports of extra-thyroidal TSHR expression and function. Aberrant TSHR expression has been documented in the orbital fibroblasts of patients with Graves’ disease in the context of thyroid associated ophthalmopathy, and TSHR-targeted treatments have shown clinical responses.(*47–51*) However, thyroid cancers have not been associated with thyroid associated ophthalmopathy. As such, it is not probable that orbital/ocular toxicities will arise with TSHR CART cell therapy. We also have not oberved any orbital toxicity in our mouse models. In addition, K1-70, the TSHR antibody from which we derived the scFv for our TSHR-CAR, was found to be well-tolerated at all dose levels in a phase I trial for patients with Graves’ disease, (*52*) and in a patient with both follicular thyroid cancer and Graves’ disease.(*53*) These clinical studies, while still nascent, demonstrate a promising safety profile. Vigilant monitoring for orbital toxicities will be incorporated into our phase I clinical trial.

CART cells targeted against ICAM-1(*54*) and GFRα4(*55*) have demonstrated efficacy in preclinical thyroid cancer models. Recently, TSHR-CART cell therapy displayed potent activity against human thyroid cancer models *in vivo.*(*38*) Furthermore, two patients with high metastatic burden and iodine-131-resistant thyroid cancer demonstrated a complete response (CR) and a near CR three months after TSHR-CART treatment. The first patient, who had papillary thyroid carcinoma, sustained CR until one year post-TSHR-CART infusion, indicating potential for long-term antitumor responses.(*56*) However, these studies focused on differentiated thyroid cancers with high TSHR expression. One patient with iodine-refractory, metastatic PDTC was treated with CART cells targeting both TSHR and CD19. This patient achieved stable disease at day 30 post-CART infusion and partial response 90 days post-CART infusion. While IHC demonstrated strong TSHR expression pre-CART infusion, it is unknown if TSHR was downregulated after the start of TSHR-CART cell therapy, leading to underwhelming therapeutic response in this dedifferentiated thyroid cancer setting.(*57*)

Several reports have shown conflicting results regarding the potential impacts of MAPK inhibitors on T cell function. One study found that dabrafenib did not suppress human T cell function, but trametinib led to an incomplete and transient inhibition of T cell proliferation and cytokine production. The addition of dabrafenib to trametinib was found to offset the negative effects of trametinib on T cell function *in vitro.* Further, trametinib did not appear to have detrimental effects on T cells, but rather increased tumor infiltration of T cells *in vivo*.(*58*) Another study found that BRAFi also enhanced tumor infiltration of adoptively transferred T cells in mice.(*59*) Other reports showed that MEKi led to reprogramming of CD8 T cells to stem cell memory phenotypes and thereby enhanced antitumor activity.(*60*) Another group reported that MEKi during chronic T cell stimulation ameliorated terminal T cell exhaustion by slowing the transcription rates of effector-and terminal-exhaustion-related genes, reducing nutrient uptake and mitochondrial ROS accumulation, and ultimately enhancing the persistence of tumor-infiltrating lymphocytes *in vivo*.(*61*) An early report of the impacts on MAPK inhibitors on CART cells showed that trametinib plus dabrafenib across a range of concentrations inhibited GD2-targeted patient-derived CART cell activation, cytokine production, and cytotoxicity; however, the authors suggested that strong stimulation of the CAR with GD2^high^ tumor cells had the potential to overcome the detrimental effects of MAPK inhibitors on CART functions.(*62*) Another GD2-CART study confirmed that trametinib inhibited CART cytotoxic abilities and cytokine secretion *in vitro*, but *in vivo*, trametinib in combination with GD2-CART resulted in enhanced tumor suppression compared to monotherapy treatment groups.(*63*) Another group showed that, while the combination of trametinib and dabrafenib did negatively impact CSPG4-targeted CART cell *in vitro* functions, these effects were milder than those seen with other MAPK inhibitor combinations.(*64*) Recently, it was reported that MEKi reduced CART cell activation, exhaustion, apoptosis, and terminal differentiation across multiple different CAR constructs in both solid and liquid tumors through downregulation of c-Fos and JunB.(*65*) While consensus has yet to be reached, more recent studies demonstrate that potential negative impacts of MAPK inhibitors on T cell function are nuanced and context-dependent, and MAPK inhibitors may actually enhance CART function in a variety of settings.

We present TSHR-CART cell therapy in combination with MAPK inhibition as a sensitizing modality, representing a major step forward in the development of a treatment strategy for PDTC and ATC as well as DTCs which have become refractory to radioiodine therapy. Based on these data, we are planning to launch a phase I clinical trial for TSHR-CART cell therapy as a monotherapy in patients with TSHR^high^ metastatic thyroid cancers and in combination with MAPK inhibitors in patients with TSHR^low^ metastatic thyroid cancers.

## MATERIALS AND METHODS

### Study design

This study aimed to assess the efficacy of TSHR-CART cell therapy as a monotherapy in TSHR^high^ thyroid cancers and in combination with MAPK inhibitors in TSHR^low^ thyroid cancers. First, the expression of TSHR was assessed and verified on patient clinical samples utilizing IHC and tissue microarray tools. Second, we designed and engineered human CART cells targeting TSHR (TSHR-CART), that were subjected to various *in vitro* and vivo *assays* to assess their effector functions (cytotoxicity, proliferation, degranulation and cytokine production). Third, we tested the addition of MAPK inhibitors *in vitro* to assess their upregulation of TSHR and sensitization prior to TSHR-CART treatment. Lastly, we performed *in vivo* assays to assess CART cell antitumor activity in combination with MAPK inhibitors in a concurrent or sequential fashion. For each experiment, the number of independent experiments (*n*) is specified in the figure legends. For *in vivo* assays, NGS mice were utilized, and they were randomly assigned to control and experimental groups, and the study was not blinded. Data collection was stopped at the predetermined times. Human endpoint criteria were determined when mice presented above 20% lost weight and tumor exceeded 700 mm^3^. No data was excluded from analysis. Sample sizes are indicated in the figure legends.

### Reagents

Detailed information on all reagents used in this article is listed in Supplementary Table S1.

### Cell lines

FTC-133, XTC.UC1, and THJ-529T cell lines were graciously provided by Dr. Copland’s lab in Mayo Clinic, Jacksonville, Florida. The FTC-133 cell line is derived from a differentiated follicular thyroid carcinoma with known mutations that include *FLCN, MSH6, NF1, PTEN, TERT*, and *TP53* (https://www.cellosaurus.org/CVCL_1219). The THJ-529T cell line was derived from a PDTC tumor containing *BRAF* and *TERT* mutations (https://www.cellosaurus.org/CVCL_9916).(66) XTC.UC1 is the only characterized HCC cell line and the only known cell line to continue to express endogenous TSHR *in vitro*.(*67*) All cell lines were DNA short tandem repeat (STR) validated prior to use. For indicated experiments, THJ-529 and FTC-133 cell lines were transduced with a firefly luciferase (Luc^+^) vector containing a hygromycin resistance gene (Promega, Madison, WI, USA) as well as a TSHR vector and a puromycin resistance gene (Genecopoeia Inc., Rockville, MD, USA). These cells were then cultured in selection media containing either hygromycin (1 µg/mL) or puromycin (1 µg/mL) to obtain a pure population as previously described.(*68*) FTC-133 and XTC.UC1 cell lines were cultured in D10 medium made with DMEM (Corning Inc., Corning, NY, USA), 10% FBS (Sigma, St. Louis, MO, USA), and 1% PSG (Gibco, Gaithersburg, MD, USA). For specific *in vitro* assays, FTC-133 cell lines were transduced with different MOIs of a TSHR vector containing puromycin resistance gene (Genecopoeia Inc., Rockville, MD, USA), and TSHR absolute counts were determined by flow cytometry using absolute counting beads (CountBright™, Invitrogen, Carlsbad, CA, USA). Mycoplasma testing was performed and confirmed to be negative monthly.

### CAR construct and CART cell production

The use of recombinant DNA in the laboratory was approved by the Mayo Clinic Institutional Biosafety Committee (IBC), IBC number HIP00000252.20. Peripheral blood mononuclear cells (PBMCs) were isolated from de-identified normal donor blood apheresis cones(*69*) obtained under a Mayo Clinic IRB approved protocol, using SepMate tubes (STEMCELL Technologies, Vancouver, BC, Canada) and Lymphoprep™ solution (STEMCELL Technologies, Vancouver, BC, Canada). T cells were separated with negative selection magnetic beads using EasySep^TM^ Human T Cell Isolation Kit (STEMCELL Technologies, Vancouver, BC, Canada). Second-generation 4-1BB-costimulated TSHR CAR constructs were synthesized *de novo* (IDT, Coralville, USA) and cloned into a third-generation lentivirus under the control of the EF-1α promotor. The TSHR-targeted single chain variable fragment was derived from an autoantibody from a patient with Graves’ disease, clone K1-70.(*70*) TSHR-CART cells were then generated through the lentiviral transduction of normal donor T cells as previously described(*71, 72*). Briefly, lentiviral particles were generated through the transient transfection of plasmid into 293T virus-producing cells (ATCC, Manassas, VA, USA) in the presence of Lipofectamine 3000 (Invitrogen, Carlsbad, CA, USA), vesicular stomatitis virus G (VSV-G), and packaging plasmids pCMVR8.74 (Addgene, Cambridge, MA, USA). The titers and multiplicity of infection (MOI) were analyzed and calculated by flow cytometry as previously described.(*68*) T cells isolated from normal donors were stimulated using anti-CD3/CD28 Dynabeads (Life Technologies, Oslo, Norway) at a 3:1 beads-to-cell ratio and then transduced 24 hours after stimulation with lentivirus particles at a MOI of 3. T cells were cultured in T cell medium containing X-VIVO™ 15 media (Lonza, Basel, Switzerland), 10% human AB serum (Corning Inc., Corning, NY, USA), and 1% PSG (Gibco, Gaithersburg, MD, USA). Magnetic bead removal and the evaluation of CAR expression on T cells by flow cytometry were performed on day 6 by staining with a goat anti-human IgG H+L secondary antibody (Invitrogen, Carlsbad, CA, USA). CART cells were harvested and cryopreserved on day 8 for future experiments. Prior to cryopreservation, CART cells were stained and evaluated to assess T-cell subset and phenotype expression patterns using the following antibodies: CD45 APC (BioLegend, San Diego, CA, USA), CD3 BV605 (BioLegend), CD4 PE-Cy7 (BD Biosciences, Franklin Lakes, NJ, USA), CD8 BV421 (BD Biosciences), CCR7 PE (BioLegend) and CD45RA APC-Cy7 (BioLegend). CART cells were thawed and rested in T cell medium overnight at 37°C and 5% CO2 prior to experimental use.

### Multi-parametric flow cytometry

Samples were prepared for flow cytometry as previously described.(*68, 71–74*) In all analyses, the population of interest was gated based on forward vs side scatter characteristics, then by singlet gating, and then by live cell gating using LIVE/DEAD™ Fixable Aqua Dead Cell Stain Kit (Invitrogen, Carlsbad, CA, USA). Flow cytometry was performed on a three-laser CytoFLEX (Beckman Coulter, Chaska, MN, USA). All analyses were performed using FlowJo v10.7.1 software (Ashland, OR, USA).

### Immunohistochemistry (IHC)

IHC was performed on paraffin-embedded tissue. Thyroid tissue microarray (TMA) was made from archival samples under Mayo Clinic IRB approval with sample size for each tumor at 1.5 mm. Paraffin blocks were cut into 5 µm thick sections. Slides were deparaffinized, rehydrated with decreasing concentrations of ethanol, and finally with water. Additionally, a normal human tissue microarray was purchased (Bio SB, Santa Barbara, CA, USA), which included 23 different normal human tissue types. Antigen retrieval was performed using Antigen Retrieval Solution (pH 6 or pH 9) (Dako, Glostrup, Denmark) for at least 15 minutes in an autoclave, and cooling for 30 minutes at room temperature. Slides were washed with 3% hydrogen peroxide (Fisher Scientific, Waltham, MA, USA) for 10 minutes to block the activity of endogenous peroxidases. Primary anti-human TSHR antibody was used for 60 minutes at room temperature at 1:500 (abcam, ab218108, Cambridge, UK). The infiltration of human T lymphocytes into tumors was assessed using anti-human CD3 at 1:500 (abcam, ab52959, Cambridge, UK). The primary antibody was detected using the Envision Labeled Kit (Dako, Glostrup, Denmark) according to the manufacturer’s protocol. After washing, slides were counterstained with Mayer’s hematoxylin (Sigma-Aldrich, St. Louis, MO, USA). Slides were dehydrated through ethanol and xylene and cover-slipped using a xylene-based mounting medium (Fisher Scientific). Samples were examined under bright-field illumination at 20x objective, and digital images were obtained using Aperio AT2 (Leica, Wetzlar, Germany). Results were processed using Aperio eSlide Manager and H-Score values were estimated using the Aperio ImageScope Software (both Aperio Technologies, Vista, CA, USA). The staining of the TMA punches was scored using an algorithm in the Imagescope Software (Aperio Technologies) created by a histologist based upon signal intensity (0, 1C, 2C, 3C). H score was then calculated based upon signal intensity and percentage: HZ(1C%!1)(2C%!2)(3C%!3). Cases were excluded from the study if a section could not be assigned a score due to insufficient quantity of tumor tissue present. Normal tissues were used as a positive and negative control.

For cell line preparation for IHC, cells were washed three times with PBS (Corning, NY, USA), scraped from the bottom of the plate using a cell lifter (Fisherbrand, Waltham, MA, USA), transferred to 50 mL polypropylene tubes and centrifuged at room temperature for 2 minutes at 500 x g. The supernatant was aspirated, and 10% neutral buffered formalin (Fisher Scientific) was added for 30 minutes at room temperature. HistoGel (Thermo Scientific) was heated in the microwave for 3 seconds at maximal power and converted to a liquid state. Cells were transferred to the HistoScreen Tissue Cassettes (Thermo Scientific) and covered with liquid Histogel. Samples were placed at room temperature and allowed to solidify. HistoGel blocks were transferred into tissue embedding cassettes (Thomas Scientific, Swedesboro, NJ, USA), dehydrated in increasing concentrations of ethanol, xylene, and paraffin embedding. Sections were prepared as described in the immunohistochemistry section. Primary anti-human TSHR antibody was used for 60 minutes at room temperature at 1:1500 (abcam, ab218108, Cambridge, UK). Envision labeled polymer (Dako, Glostrup, Denmark) was used for 30 minutes as a secondary antibody. Slides were stained with diaminobenzidine tetrahydrochloride (DAB) chromogen (Dako) for 5 minutes at room temperature and counterstained with Mayer’s hematoxylin (Sigma-Aldrich, St. Louis, MO, USA). Control staining with hematoxylin and eosin (Sigma-Aldrich) was also performed. Aperio AT2 scanner (Leica, Wetzlar, Germany) was used for the digital image, at 20x objective. Images were analyzed using the Aperio ImageScope software (Aperio Technologies).

### Immunocytochemistry (ICC)

THJ-529 cells (2 x 10^4^ cells/chamber) were plated in a 4-well chamber slide (Nunc, Thermo Scientific, Waltham, MA, USA), incubated overnight. After 24 hours, media was aspirated, and cells were washed three times with PBS (Corning, NY, USA). Then, 2% paraformaldehyde was added for 20 minutes at room temperature. Again, cells were washed three times with PBS. Next, ice-cold 100% methanol (Fisher Scientific) was added for 7 minutes at -20°C. Methanol was aspirated, and samples were allowed to air dry for 10 minutes. Next, serum-free-blocking diluent (Dako) was used for 30 minutes at room temperature. Primary anti-human TSHR antibody was used for 60 minutes at room temperature at 1:1500 (abcam, ab218108, Cambridge, UK). Slides were subsequently prepared as described in the immunohistochemistry section.

### *In vitro* T cell function assays

Cytotoxicity assays were performed as previously described.(*68, 71–74*) Wildtype (TSHR^-^) Luc^+^ FTC-133, TSHR^+^ Luc^+^ FTC-133, wildtype (TSHR^-^) Luc^+^ THJ-529, or TSHR^+^ Luc^+^ THJ-529 cells were used as target cells. UTD or TSHR-CART cells were co-cultured with target cells at various effector: target (E:T) ratios in T cell medium. Cytotoxicity was calculated by bioluminescence imaging on the Promega GloMax Explorer (Promega Corporation, Fitchburg, WI, USA) as a measure of residual live cells at indicated time points. Samples were treated with 1µL D-luciferin (30µg/mL) per 100µL sample volume (Gold Biotechnology, St. Louis, MO, USA) prior to imaging. Proliferation assays were performed utilizing two approaches as previously described.(*68, 71–74*) Briefly, for the first approach, TSHR-CART cells were stained with CFSE (Invitrogen) cell dye on day 0, then washed and counted prior to being plated at a 1:1 E:T ratio with target cell lines. Each assay included T cells with media only as a negative control and T cells with PMA and ionomycin as a positive control. On day 5, percent of CFSE CART cells was determined via flow cytometry after staining with APC-H7 anti-human CD3 (BD Biosciences) and Zombie R718™ (BioLegend) dye. Alternatively, the same assay was repeated but without the addition of CFSE and with LIVE/DEAD™ Fixable Aqua as the viability dye.

For *in vitro* experiments testing the cytotoxic and proliferation effects of TSHR-CART cells exposed to different levels of TSHR on target cells, TSHR-CART cells were co-cultured with FTC-133 tumor cells expressing different levels of TSHR and cytotoxicity and proliferation was measured as described above.

For cytokine assays, media was collected on day 3 of co-culture of TSHR-CART cells with target cells, and supernatants were isolated and purified via centrifugation as previously described(*71*). Supernatants were prepared following manufacturer’s protocol for the MILLIPLEX® Human Cytokine/Chemokine/Growth Factor Panel A 38 Plex Premixed Magnetic Bead Panel (Millipore Sigma, Ontario, Canada). Cytokine data was collected using a Luminex LX200 (Millipore Sigma). For degranulation assays (*68, 71*), UTD or TSHR-CART cells were incubated with various target cells at an E:T ratio of 1:5, and antibodies against CD107a FITC (BD Biosciences), CD28 (BD Biosciences), and CD49d (BD Biosciences), as well as monensin (BioLegend), were added at the start of the incubation. After 4 hours, cells were harvested and stained with LIVE/DEAD™ Fixable Aqua. Cells were then fixed and permeabilized (FIX & PERM Cell Fixation & Cell Permeabilization Kit, Life Technologies, Oslo, Norway) and stained for CD3 APC (, eBioscience, San Diego, CA, USA) and intracellular cytokines including IL-2 PE CF594 (BD Horizon), GM-SCF BV421 (BD Horizon) and MIP-1β Pe-Cy7 (BD Pharmigen).

For *in vitro* assays performed to assess the impact of MAPK inhibitors on TSHR-CART cells, TSHR-CART cells were cultured with target tumor cells or media alone, along with either vehicle control (DMSO), trametinib (10nM) alone, or trametinib plus dabrafenib (10nM). Effector functions of TSHR-CART cells (proliferation, cytotoxicity, degranulation and cytokine profile) were assessed as described above. Alternatively, TSHR-CART cells were cocultured in an *in vitro* model for CART cell exhaustion previously described by our laboratory.(*40*) Here, TSHR-CART cells were cultured at a 1:1 ratio with irradiated target cells along with either vehicle control (DMSO), trametinib (10nM) alone, or trametinib plus dabrafenib (10nM). Drug-containing media and irradiated target cells were added to the TSHR-CART cell cocultures every other day for 7 days. Then, chronically stimulated TSHR-CART cells were isolated using CD4 (Miltenyi Biotec, Auburn, CA, USA) and CD8 (Miltenyi Biotec) microbeads and were subsequently plated with fresh, non-irradiated target cells to assess cytotoxic and proliferative capacities as described above.

### In Vivo Models

Male and female 6-8-week-old NSG mice were purchased (Jackson Laboratories, Bar Harbor, ME, USA) and maintained in an animal barrier space that is approved by the institutional Biosafety Committee for BSL2+ level experiments at the Mayo Clinic animal facility (IBC #HIP00000252.20). Mice were treated on an IACUC-approved protocol (A00001767).

#### XCT.UC1 model

The purpose of this model was to test initial TSHR-CART activity in a thyroid cancer model which retains endogenous TSHR expression *in vivo*. XCT.UC1 is a Hurtle cell carcinoma cell line which retains TSHR expression when passaged in mice but not *in vitro*. 1 x 10^7^ XTC.UC1 cells were suspended in a 50% Matrigel basement medium and subcutaneously injected into the rear flank of NSG mice. Serial caliper measurements were performed to confirm tumor engraftment. Mice were then randomized based on tumor burden to receive 1 x 10^7^ UTD or TSHR-CART. Tumor growth was monitored by serial caliper measurements (width x height x length x 0.523 = mm^3^). Mice were euthanized once IACUC-approved endpoint criteria were met, and remaining tumors were collected for IHC as indicated in each specific experiment.

#### TSHR^+^ THJ-529 model

The purpose of this model was to assess a TSHR-CART dose escalation. TSHR^+^ THJ-529 is a poorly differentiated thyroid cancer cell line made to overexpress TSHR. 2.5 x 10^6^ TSHR^+^ THJ-529 cells were suspended in a 50% Matrigel basement medium and subcutaneously injected into the rear flank of NSG mice. Serial caliper measurements were performed to confirm tumor engraftment. Mice were then randomized based on tumor burden to receive 5 x 10^6^ UTD, 5 x 10^6^ TSHR-CART, or 1 x 10^7^ TSHR-CART. Tumor growth was monitored by serial caliper measurements (width x height x length x 0.523 = mm^3^). Mice were euthanized once IACUC-approved endpoint criteria were met, and remaining tumors were collected for IHC as indicated in each specific experiment.

#### TH-560 model

The purpose of this model was to assess TSHR upregulation or TSHR-CART activity after MAPKi administration. This model reflects the strategy of TSHR-CART and MAPK inhibitor combination to be used in patients with ATC. TH-560 is an ATC PDX with downregulated TSHR expression. All NSG mice were surgically engrafted with 5mm^3^ ATC PDX TH-560 tumors in the rear flank. In TSHR upregulation experiments, mice were treated with the indicated doses of MEKi/BRAFi (trametinib, dabrafenib, or R05126766). Upon euthanasia, tumors were harvested for IHC.

In the MAPK inhibitor and TSHR-CART combination experiments, upon tumor engraftment, mice were treated with a sub-therapeutic dose of 1mg/kg trametinib and 1.5mg/kg dabrafenib as a sensitizing regimen to induce TSHR expression on ATC tumor cells. On Day 7, additional mice were surgically engrafted with 5mm^3^ ATC PDX tumors in the same manner as the first arm but were treated with placebo rather than MAPK inhibitors. In the first MAPK inhibitor and TSHR-CART combination experiment, in which mice were treated sequentially or concurrently with MEKi/BRAFi and TSHR-CART cells, all mice were randomized based on tumor burden when tumors reached ∼100mm^3^ and injected with 2 x 10^7^ UTD or TSHR-CART; this higher TSHR-CART dose was a further dose escalation experiment. In the second MAPK inhibitor and TSHR-CART combination experiment, in which mice were treated concurrently with MEKi/BRAFi and TSHR-CART cells with various MEKi/BRAFi treatment stopping points, all mice were randomized based on tumor burden when tumors reached ∼100mm^3^ and injected with 1 x 10^7^ UTD or TSHR-CART. This lower dose of TSHR-CART was used since no further treatment benefit was observed with the higher 2 x 10^7^ dose. For all mice in the MAPK inhibitor and TSHR-CART combination experiments, tumor growth was monitored by serial caliper measurements (width x height x length x 0.523 = mm^3^). Weekly tail vein bleeding after injection of CART cells was performed to assess T cell expansion. Mouse peripheral blood was lysed using BD FACS Lyse (BD Biosciences) and then analyzed with flow cytometry as previously described.(*68*) Mice were euthanized at IACUC-approved endpoints.

### Statistical analysis

GraphPad Prism (La Jolla, CA, USA) and Microsoft Excel (Microsoft, Redmond, WA, USA) were used to analyze raw experimental data. Statistical tests are described in figure legends.

## Author Contributions

CMR, ELS, and JJG wrote, reviewed, and edited the manuscript, designed and performed experiments, and analyzed data. JJG, JLM, EEM, AAD, TNH, GED, LKM, BLK, EET, RLS, ELS, MLP, AAD, CMS, IC, OLS, KY, JMF, OLGR, HX, MHT, KJS, and EJO aided in data collection. GO aided in patent and intellectual property management. YQ, HWT, RCS, and AZ aided in data collection and troubleshooting. JAC and SSK formulated the initial concept, designed experiments, supervised the study, and wrote the manuscript. All authors edited and approved the final version of the manuscript.

## Acknowledgements

This study was partly funded through the following:

Mayo Clinic Center for Regenerative Biotherapeutics (JAC and SSK)

Mayo Clinic Comprehensive Cancer Center (SSK)

National Institutes of Health R37CA266344 and R01AI79974 (SSK)

Department of Defense grant CA201127 (SSK)

Generous support from Renee Cousins King, M.D.

---

**Sup. Fig. S1.**
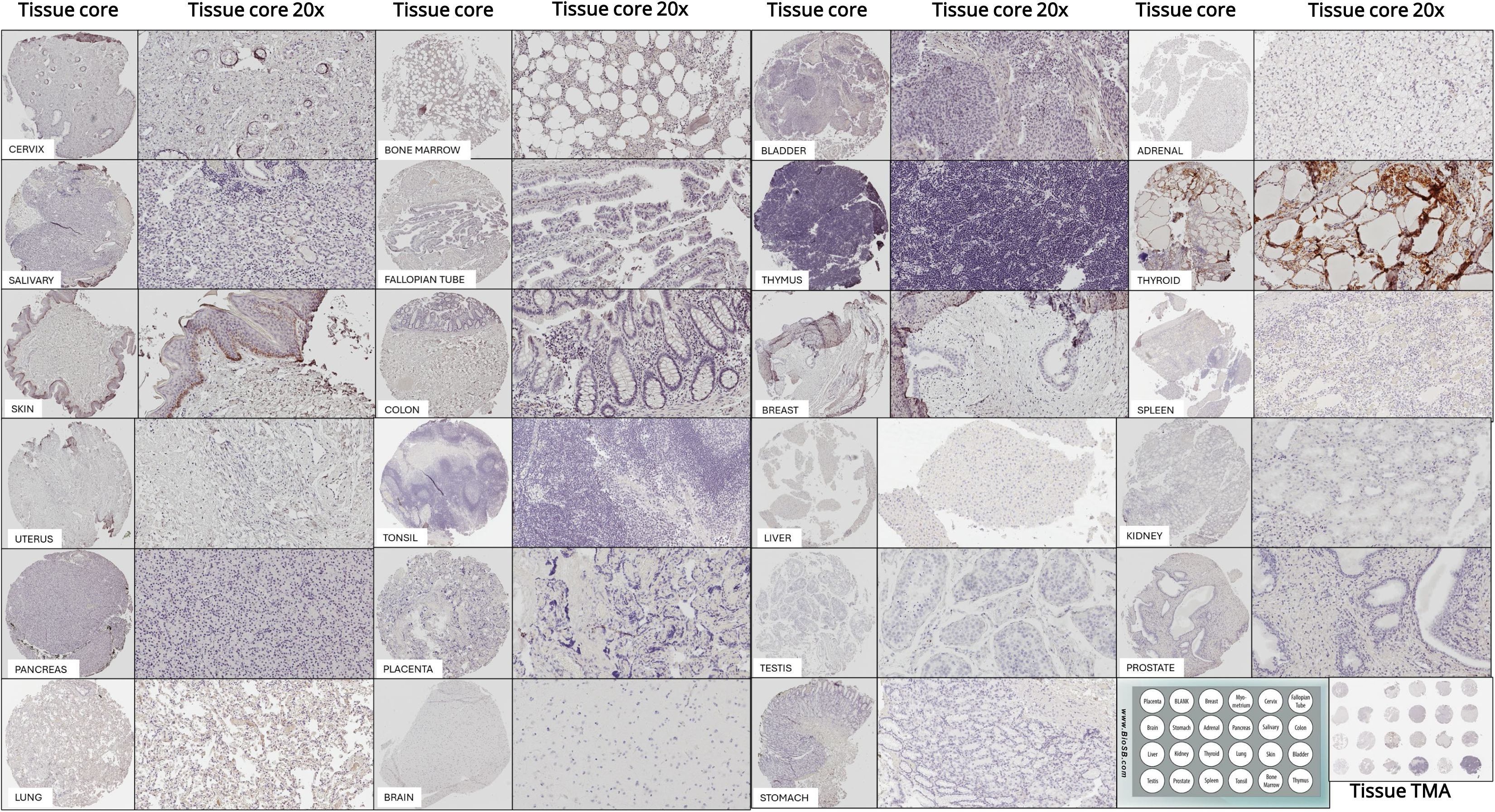

**Sup. Fig. S2.**
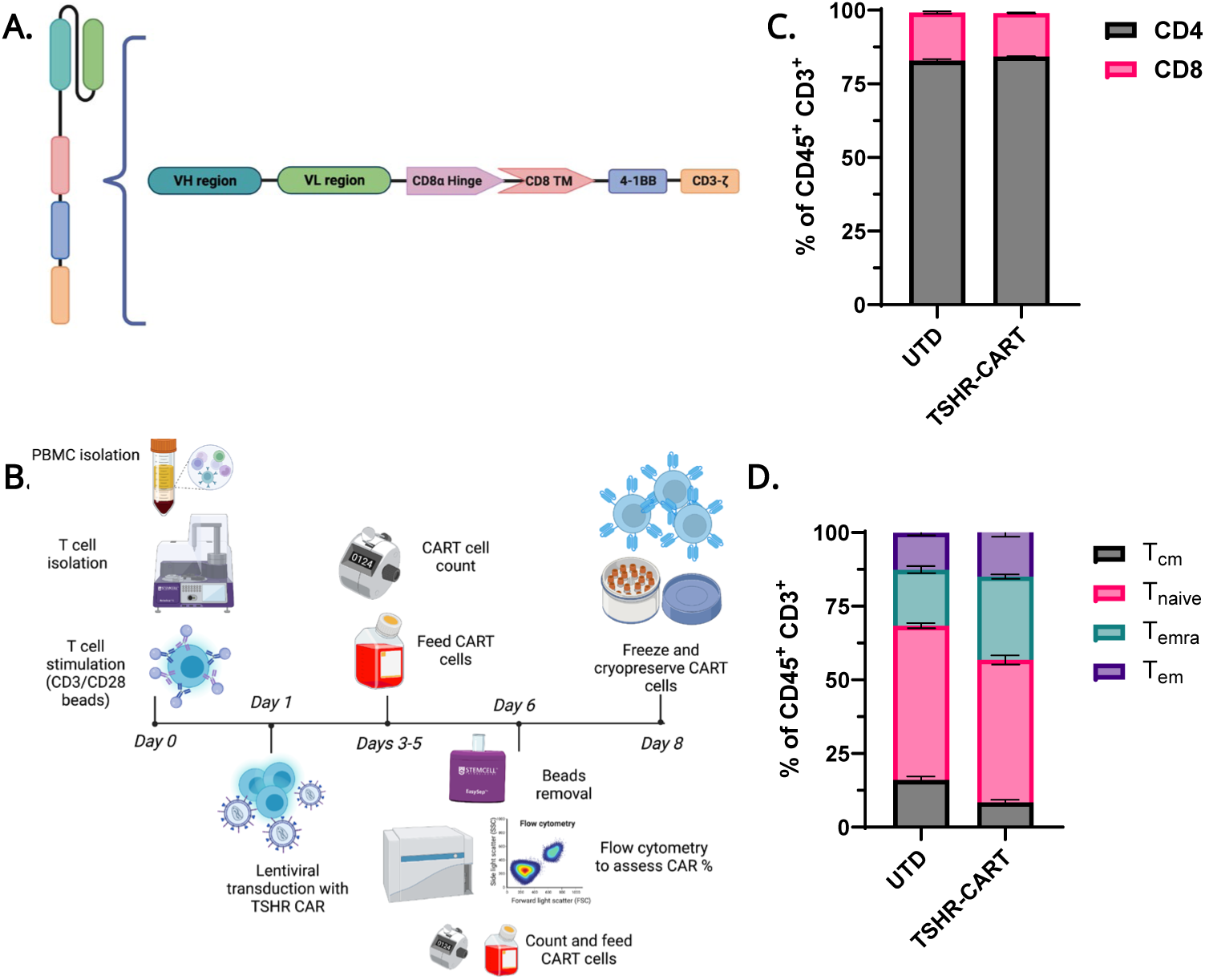

**Sup. Fig. S3.**
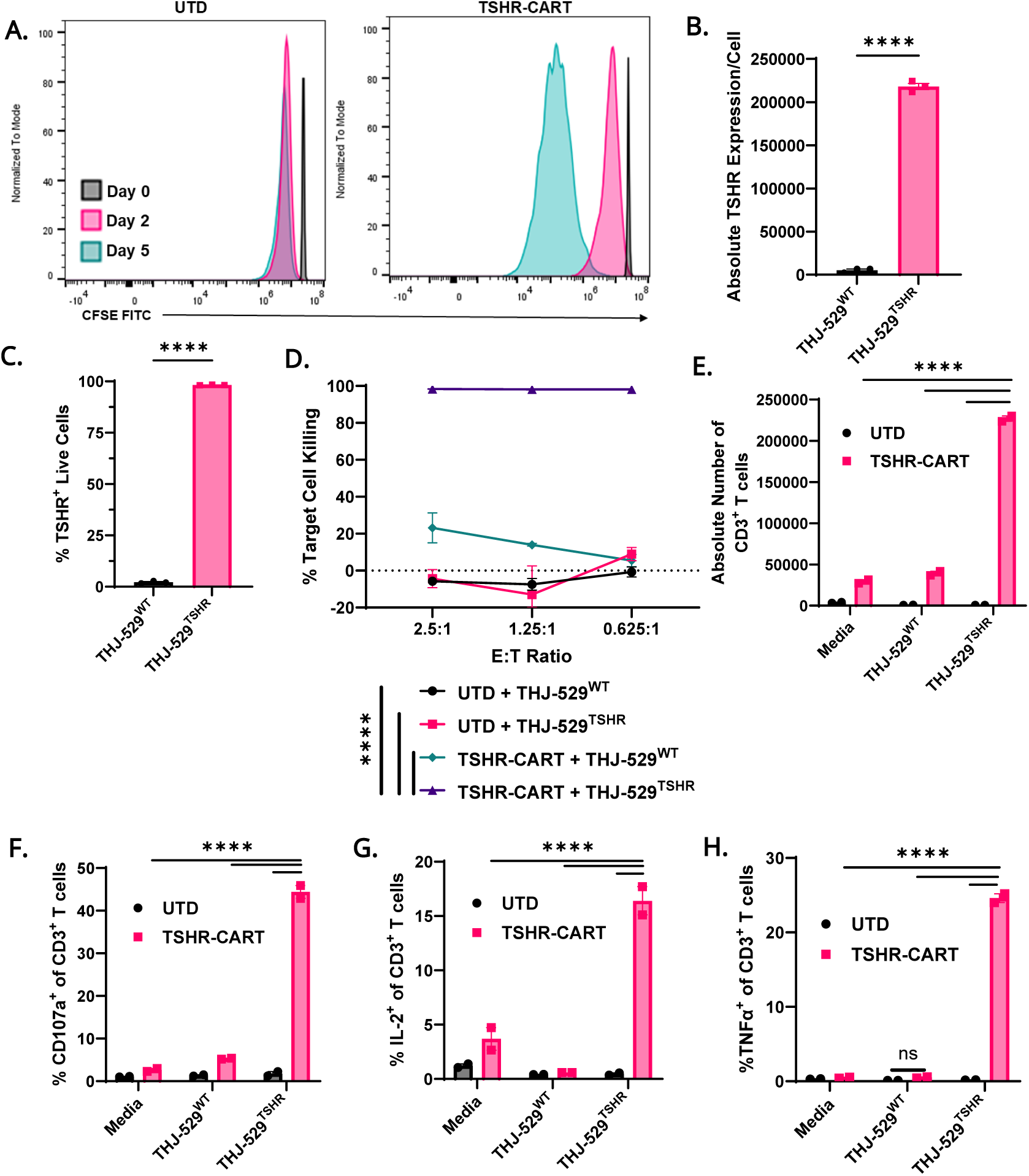

**Sup. Fig. S4.**
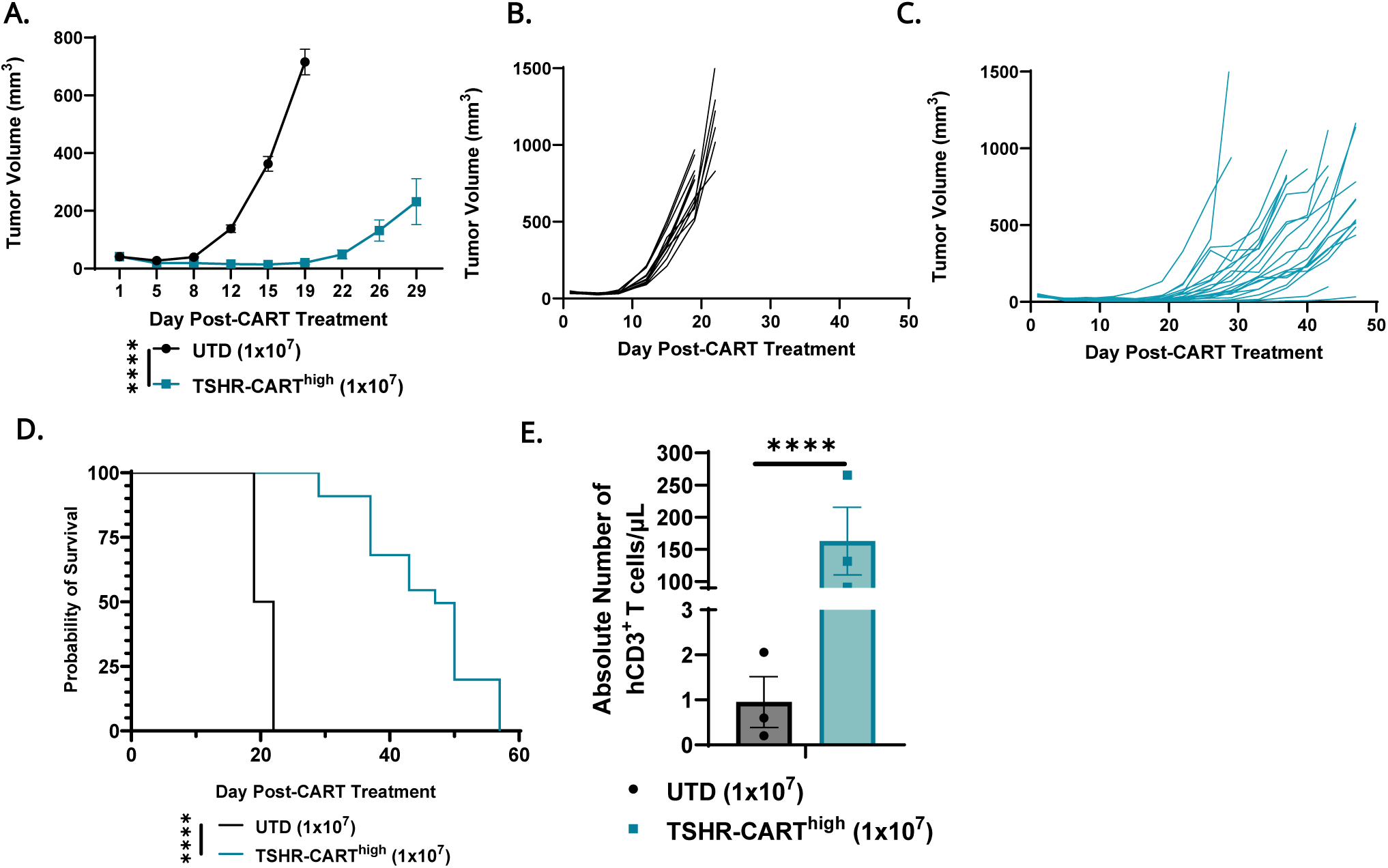

**Sup. Fig. S5.**
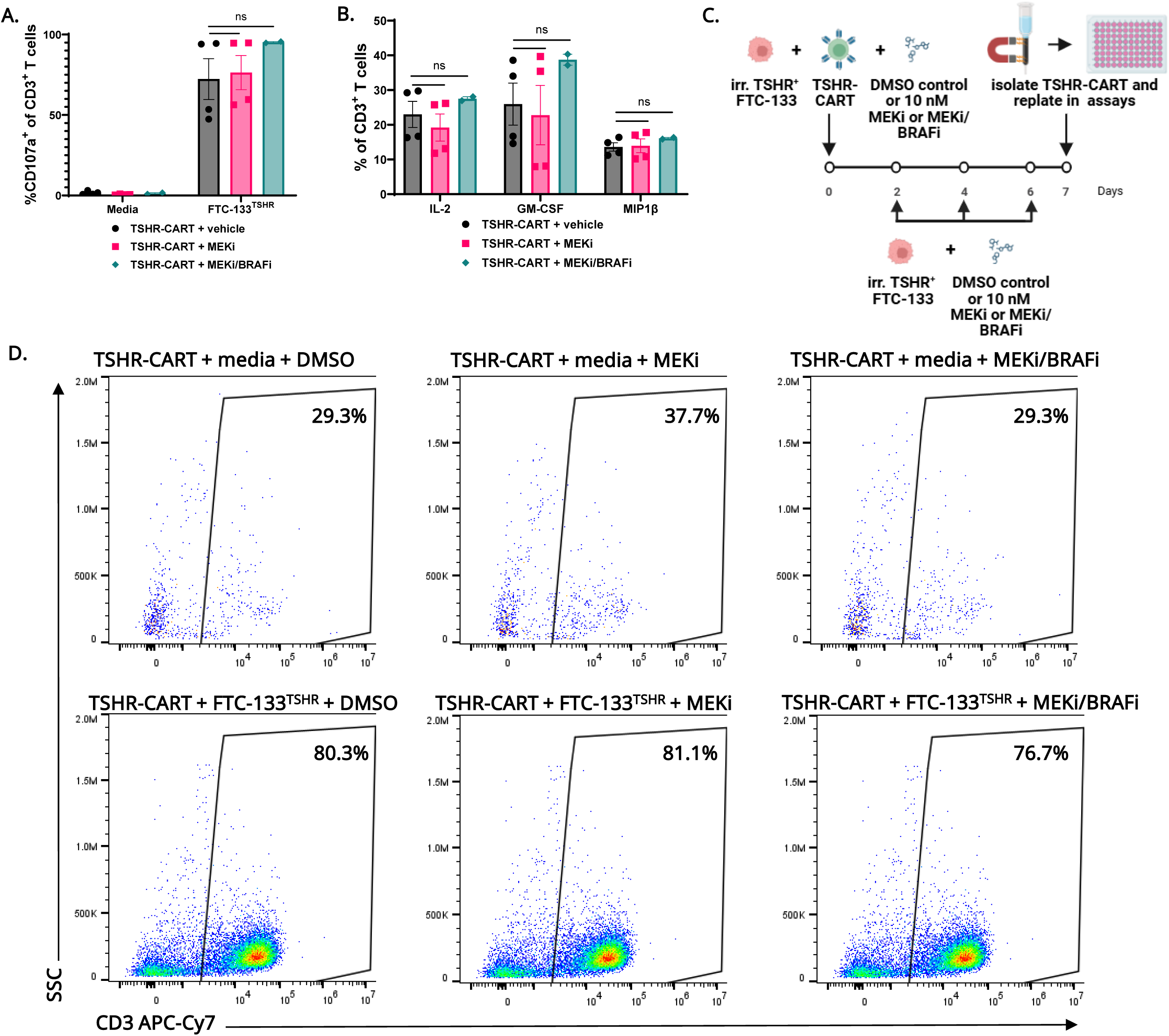

**Sup. Fig. S6.**
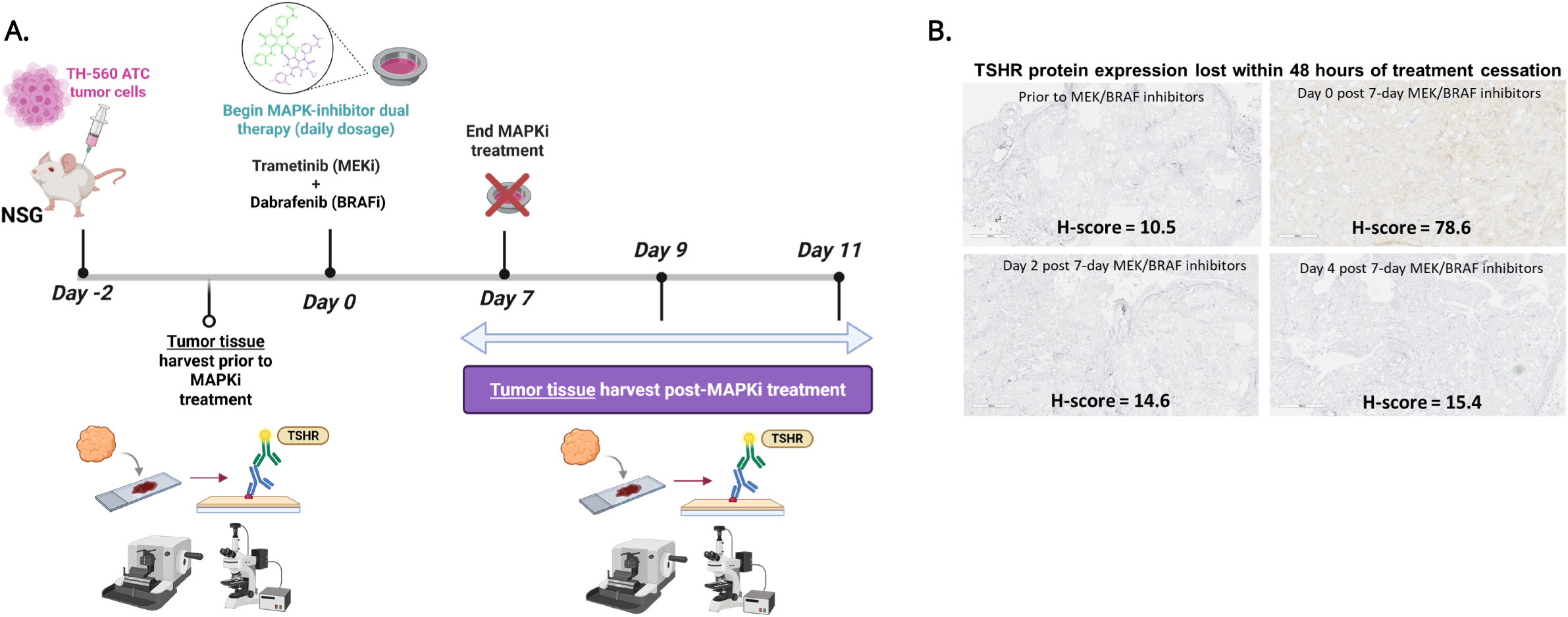

**Sup. Fig. S7.**
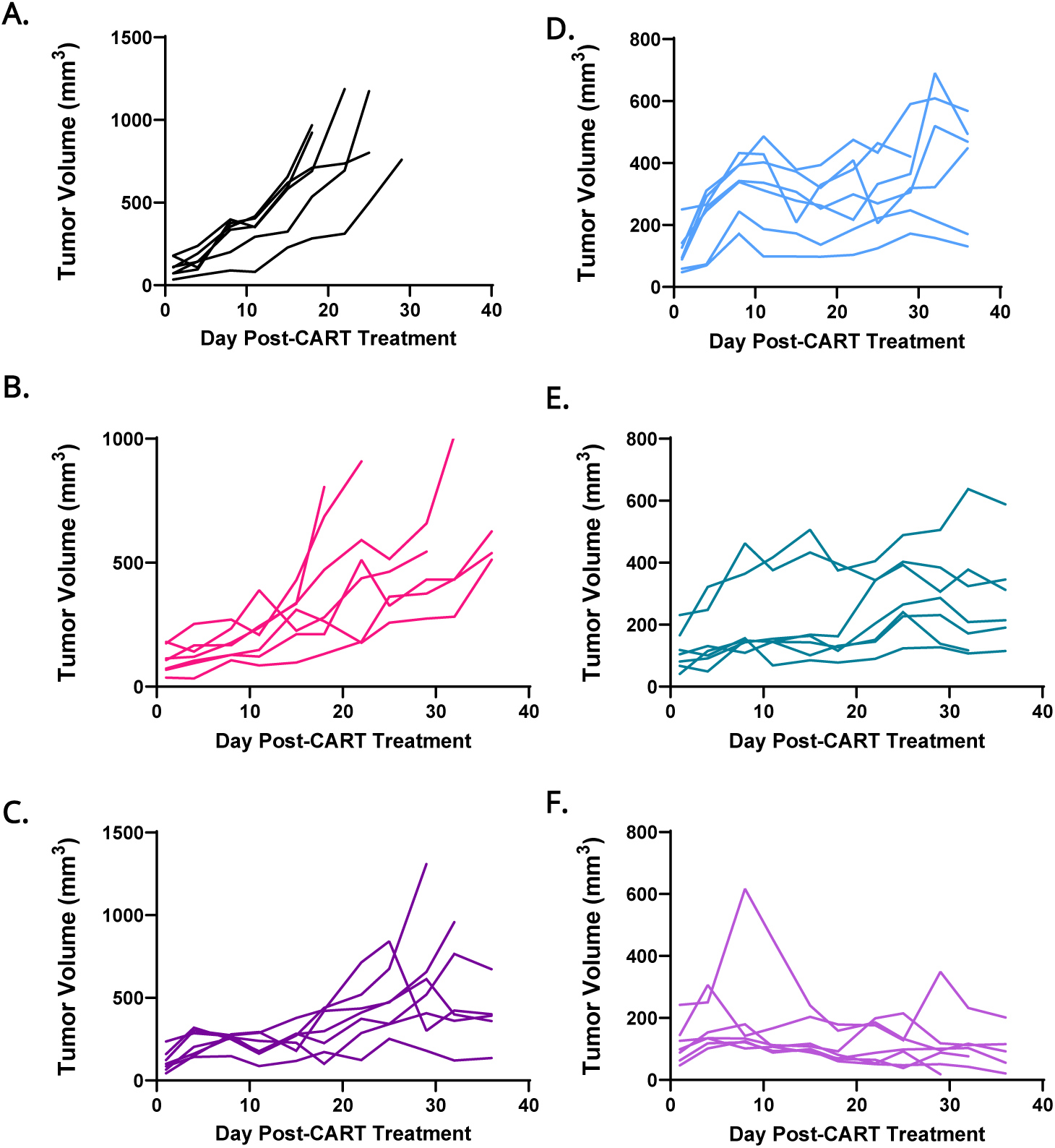

**Sup. Fig. S8.**
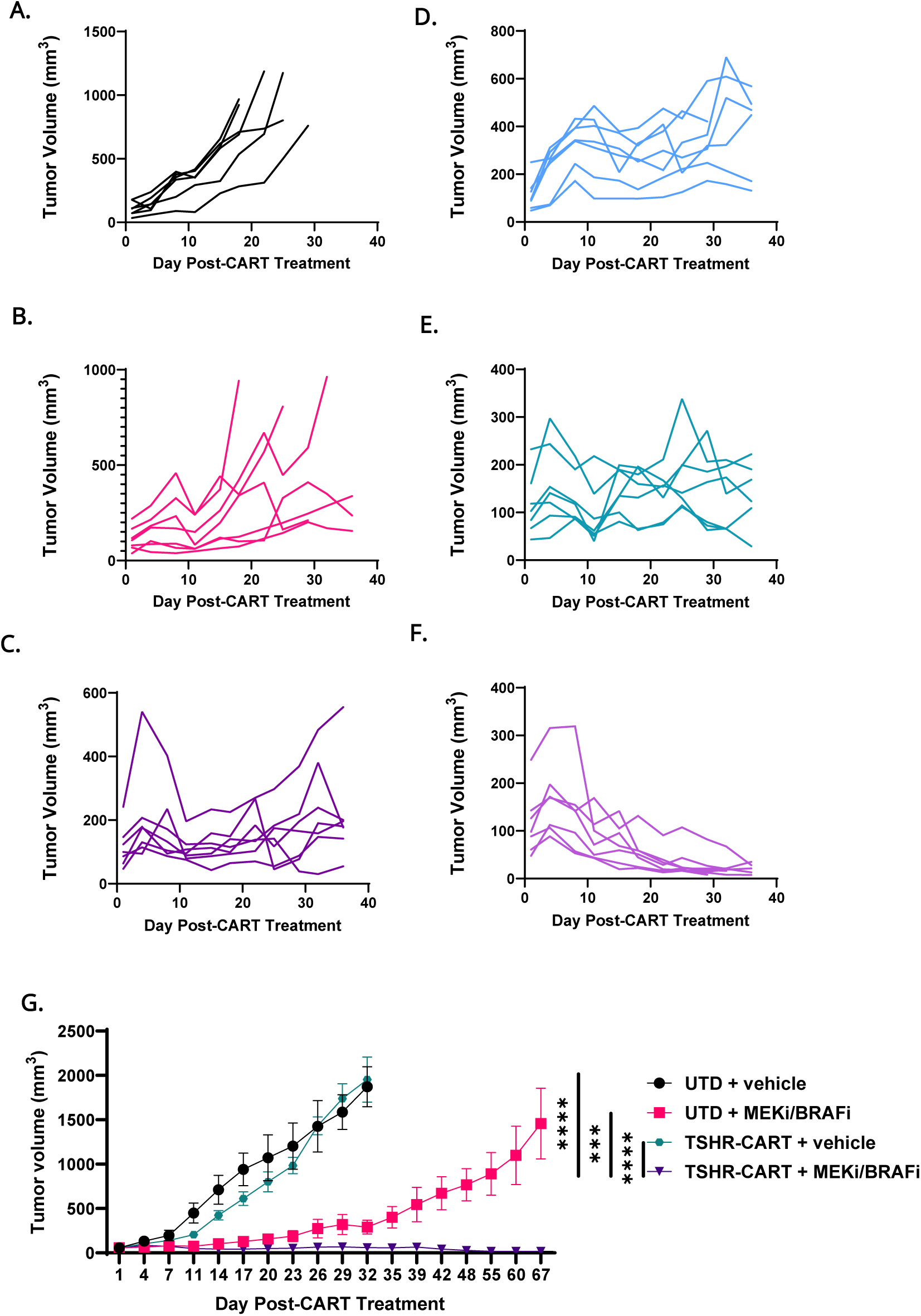

**Sup. Fig. S9.**
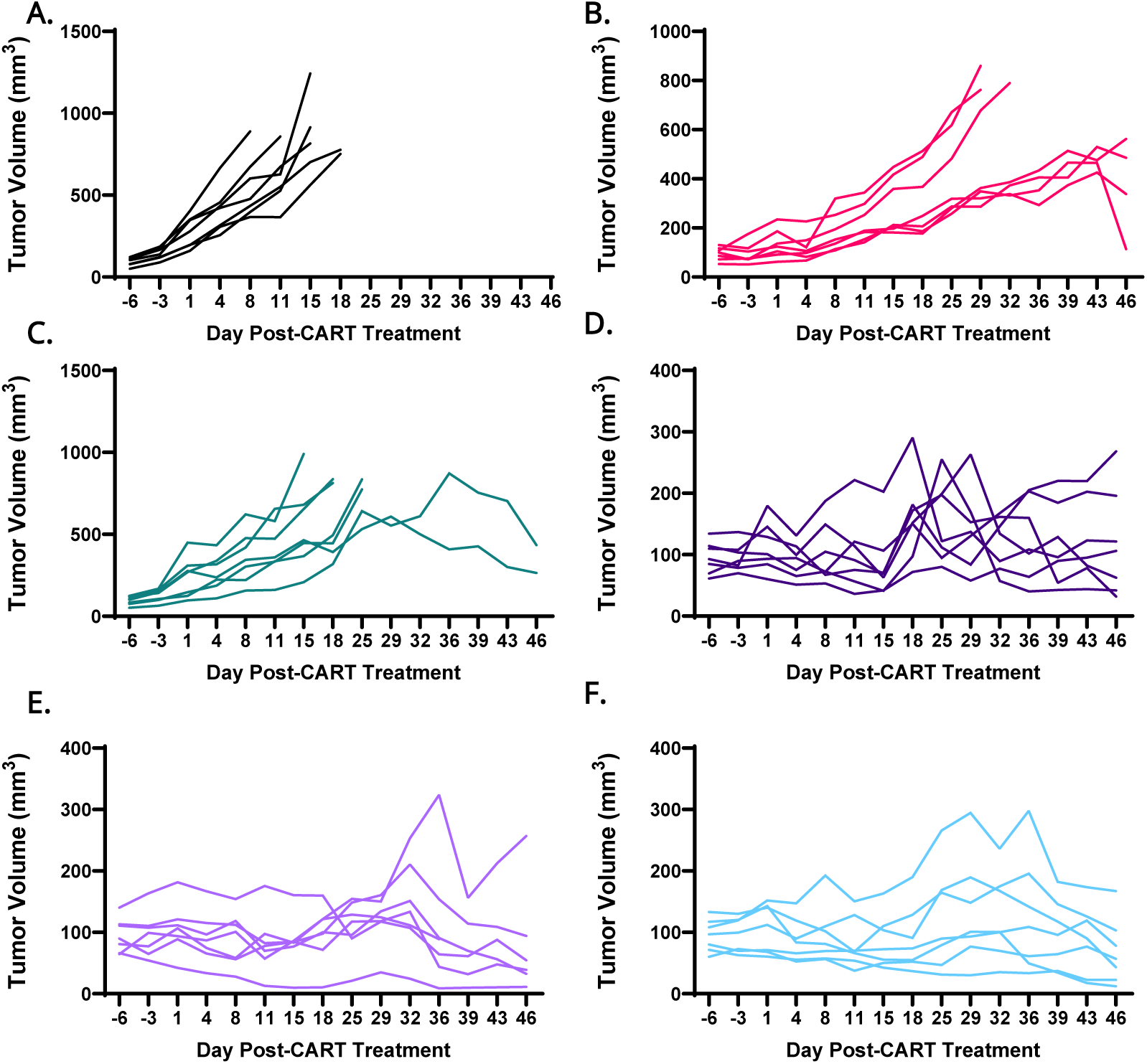

