## Supplemental for "Optimized CART Cell Therapy for Metastatic Aggressive Thyroid Cancer"

Sup. Fig. S1

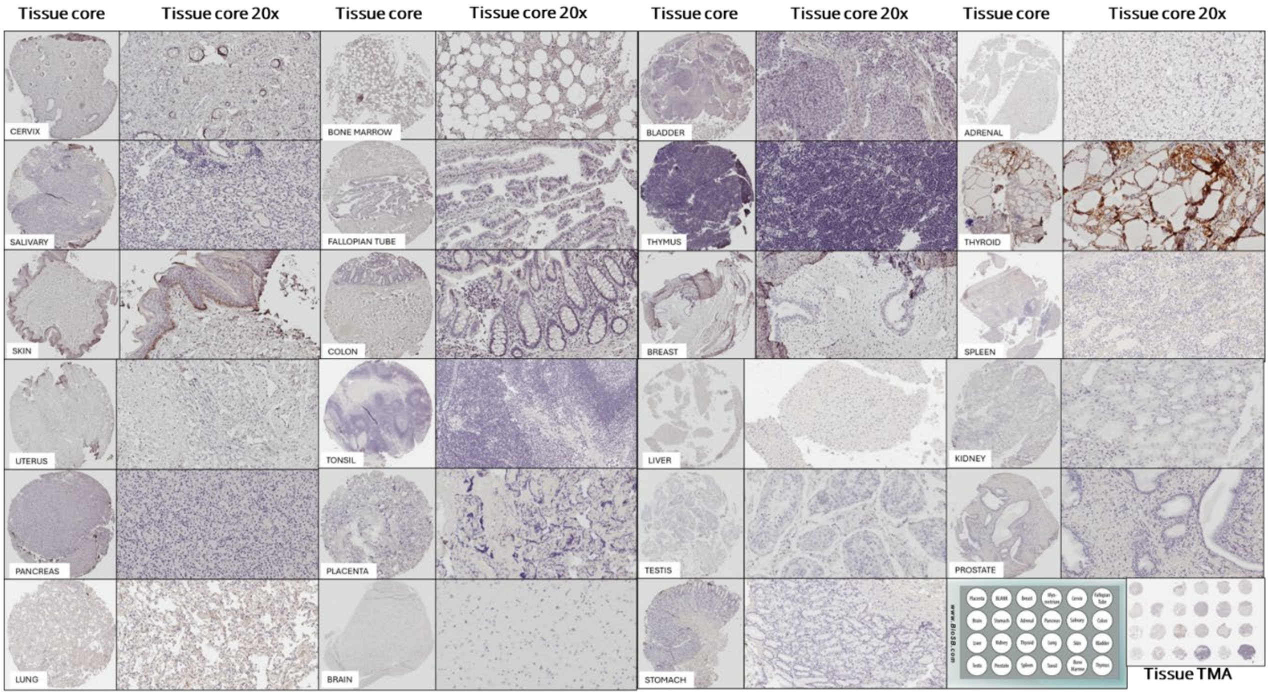


**Supplementary Figure S1. Normal human tissue microarray from 23 different human normal tissues stained with TSHR.** Tissue columns showing the TMA of 23 different tissue organs (placenta, brain, liver, testis, stomach, kidney, prostate, breast, adrenal, spleen, myometrium (uterus), pancreas, lung, tonsil, cervix, salivary, skin, bone marrow, fallopian tube, colon, bladder, thyroid, and thymus). Each column represents (1) whole tissue core, (2), 20x magnification and (3) slide area from where the presented image was taken.

Sup. Fig. S2

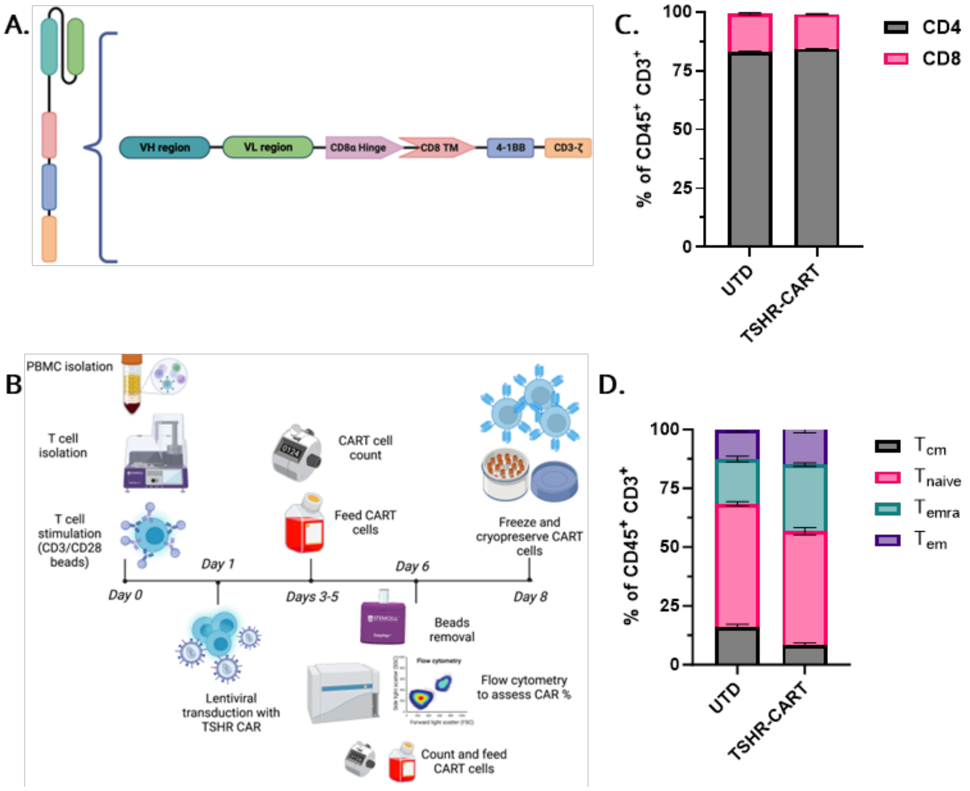


**Supplementary Figure S2. TSHR-CAR design, manufacturing, and phenotype. A)** TSHR-CAR schematic. The CAR was composed of anti-TSHR scFv (derived from anti-TSHR human autoantibody clone K1-70), CD8α hinge and transmembrane domain, 4-1BB costimulatory domain, and CD3ζ signaling domain. **B)** TSHR-CART production schematic. T cells were isolated from healthy donor PBMCs and stimulated with CD3/CD28 beads on Day 0. On Day 1, T cells were lentivirally transduced with CAR transgene. TSHR-CART cells were expanded in T cell media. On Day 6, CD3/CD28 beads were removed via magnetic separation, and T cells were assessed for CAR expression by flow cytometry. On Day 8, TSHR-CART cells were counted and cryopreserved for future use. **C)** Percentages of CD4$^+$ and CD8$^+$ T cell subsets were assessed by flow cytometry on Day 8 (1 representative biological replicate shown). **D)** T cell phenotypes were assessed by flow cytometry of CD45RA and CCR7, which are associated with the following phenotypes: central memory ($T_{cm}$ CCR7$^+$ CD45RA$^-$), naïve ($T_{naive}$, CCR7$^+$ CD45RA$^+$), terminal effector ($T_{emra}$ CCR7$^-$ CD45RA$^+$) and effector memory ($T_{em}$ CCR7$^-$CD45RA$^-$)[1 representative biological replicate shown].

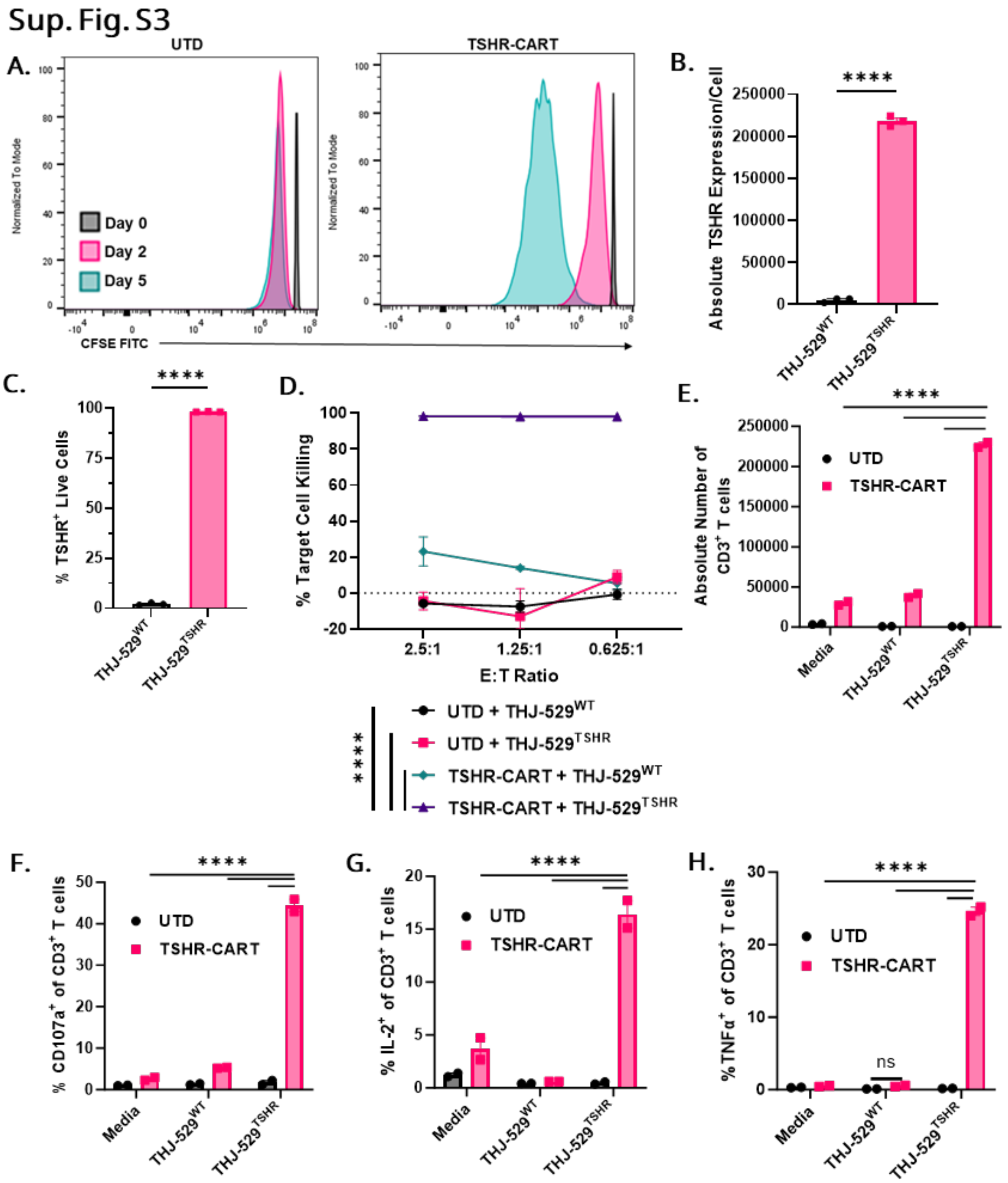


**Supplementary Figure S3. TSHR-CART cells demonstrate potent antigen-specific anti-tumor activity *in vitro*. A)** Histogram demonstrating the proliferative capacity, via CFSE staining at different time points, of TSHR-CART cells vs UTD cells. **B-C)** The PDTC cell line THJ-529 was transduced to overexpress TSHR. TSHR absolute counts or percentage of positive cells of wildtype or $TSHR^+$ THJ-529 was assessed by flow cytometry (mean and SEM; ****$p < 0.0001$, unpaired two-tailed t-test; 3 technical replicates). **D)** Luciferase$^{+}$ TSHR$^{+}$ or TSHR$^{-}$ THJ-529 target cells were co-cultured with UTD or TSHR-CART cells at varying effector-to-target ratios, and cytotoxicity was measured via bioluminescence after 48 hours (mean and SEM; ***p < 0.001, ****p < 0.0001, two-way ANOVA; 1 biological replicate). **E)** UTD or TSHR-CART cells were co-cocultured with media alone (negative control) or TSHR$^{+}$ or TSHR$^{-}$ THJ-529 target cells. After five days, T cell proliferation was assessed by flow cytometry using absolute counts of CD3$^{+}$ cells (mean and SEM; ****p < 0.0001, two-way ANOVA; 1 biological replicate). **F-H)** UTD and TSHR-CART cells were cultured with media alone or with TSHR$^{-}$ or TSHR$^{+}$ THJ-529 tumor cells at a 1:5 E:T ratio. Four hours later, cells were flowed to assess CD107a$^{+}$ (F), IL-2 (G), and TNFα (H) (mean and SEM; ****p < 0.0001; two-way ANOVA; 1 biological replicate).

Sup. Fig. S4

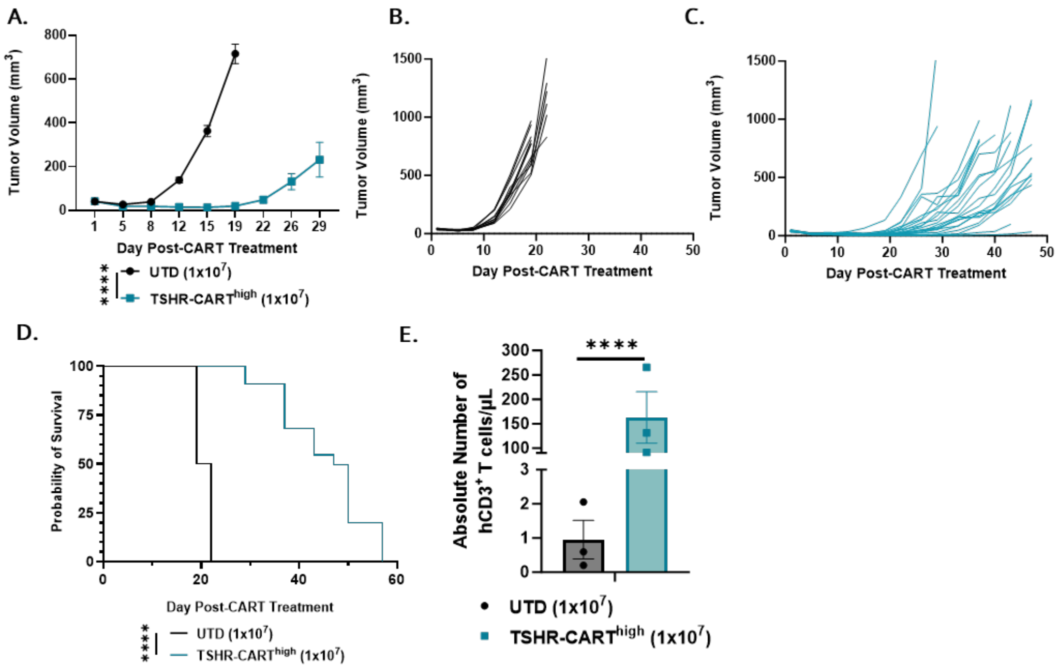


**Supplementary Figure S4. TSHR-CART cells exhibit antitumor activity in TSHR$^+$ ATC PDX models. A)** Tumor volume was assessed via serial caliper measurements (mean and SEM; ***p < 0.0001, two-way ANOVA; n=12 mice in the UTD group, n=22 mice in the TSHR-CART group). **B-C)** Individual tumor growth curves are shown for each treatment group, including UTD (**B**) and TSHR-CART$^{high}$ (**C**). N=12-22 mice/group. **D)** Kaplan-Meier survival curve (****p < 0.0001, Log-rank test; n=12 mice in the UTD group, n=22 mice in the TSHR-CART group). **E)** Peripheral blood was collected 21 days after CART administration, and T cell proliferation was assessed by flow cytometry using absolute counts of hCD3$^+$ cells (mean and SEM; ****p < 0.0001, two-tailed unpaired t-test; n = 3 mice per group).

Sup. Fig. S5

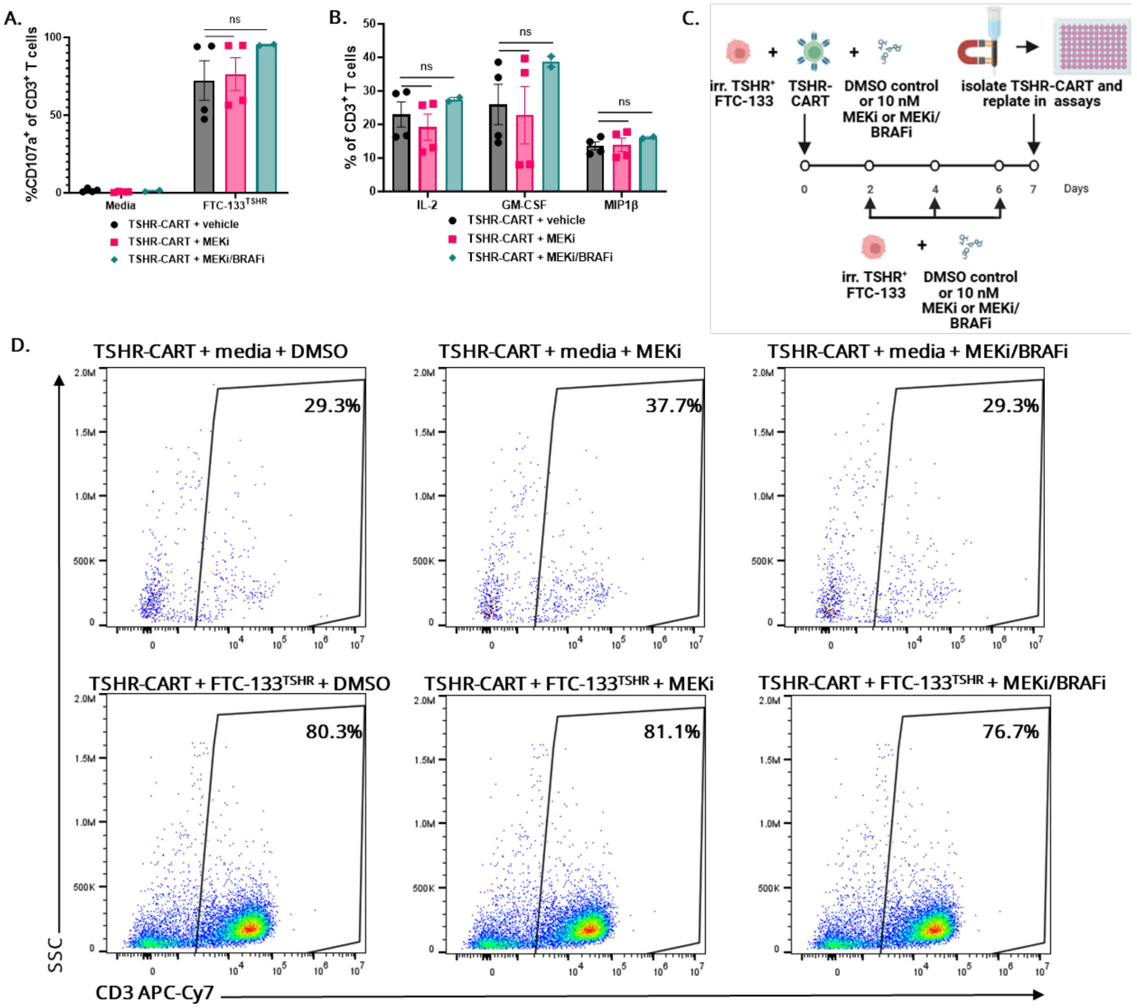


**Supplementary Figure S5. MAPK inhibitor treatment is not detrimental for TSHR-CART cell effector functions *in vitro*. A-B)** TSHR-CART cells were plated with media alone or with TSHR$^{+}$ FTC-133 tumor cells at a 1:5 E:T ratio in the presence of MEKi or combined MEKi/BRAFi. Four hours later, cells were fixed and permeabilized, and CD107$^{+}$ degranulation marker (**A**) and IL-2, GM-CSF, and MIP1β (**B**) were assessed via flow cryometric analysis (mean and SEM; ns = not significant, two-way ANOVA; 2 biological replicates). **C)** Experimental schema of repeated stimulation experiments. **D)** Representative flow plots of CD3$^{+}$ gated on live singlet cells in proliferation assays after extended MAPK inhibitor culture.

Sup. Fig. S6

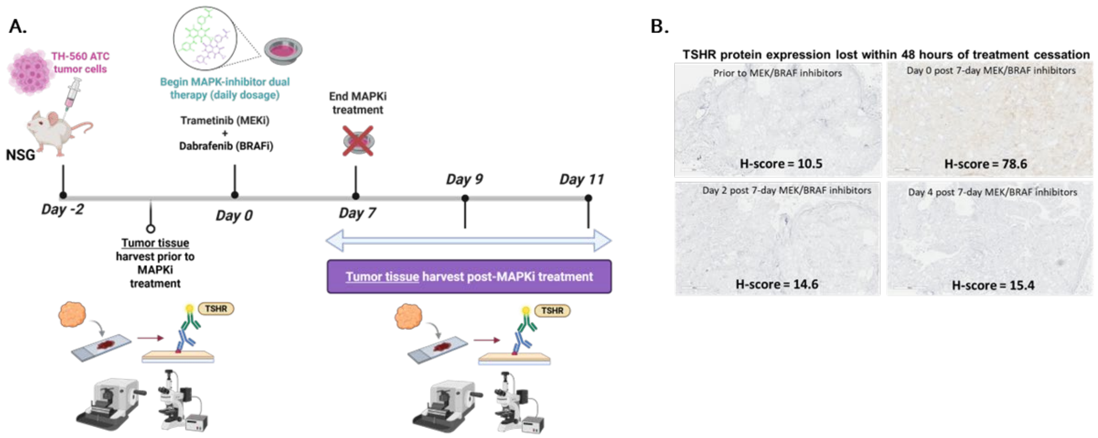


**Supplementary Figure S6. Representative IHC staining shows human TSHR expression in thyroid PDX tumors excised from mice at various timepoints. A)** Experimental schema of an ATC PDX model to assess TSHR expression. NSG mice were subcutaneously engrafted with Th-560 ATC PDX tumors. 2 days later, when tumors reached approximately 100 mm$^3$, mice were orally treated with MAPK inhibitors for seven days. Tumors were harvested and assessed for TSHR expression via IHC prior to MAPK inhibitor treatment, at day 7 of treatment and 2- and 4-days post-treatment cessation. **B)** Representative IHC staining of TSHR expression in excised tumors. H-scores provided are based on 0 – 3 IHC scoring and percentage of total areas each score (0x%0 + 1x%1 + 2x%2 + 3x%3 = H Score; 0 – 300 range with statistical analysis).

Sup. Fig. S7

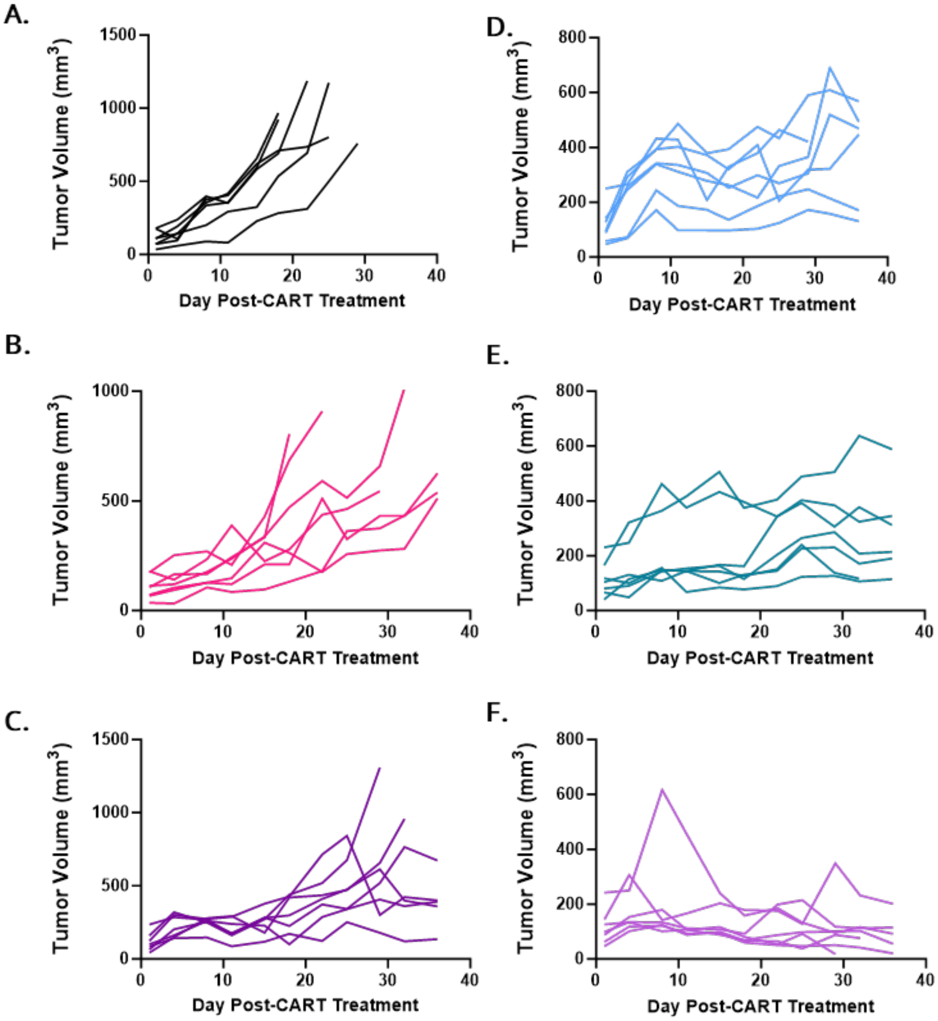


**Supplementary Figure S7. TSHR-CART cells showed the most potent antitumor activity when combined with concurrent MEKi administration. A-F)** Individual tumor growth curves are shown for each treatment group, including UTD + vehicle (**A**), sequential MEKi → UTD (**B**), concurrent MEKi + UTD (**C**), TSHR-CART + vehicle (**D**), sequential MEKi → TSHR-CART (**E**), and concurrent MEKi + TSHR-CART (**F**). N = 7 mice/group.

Sup. Fig. S8

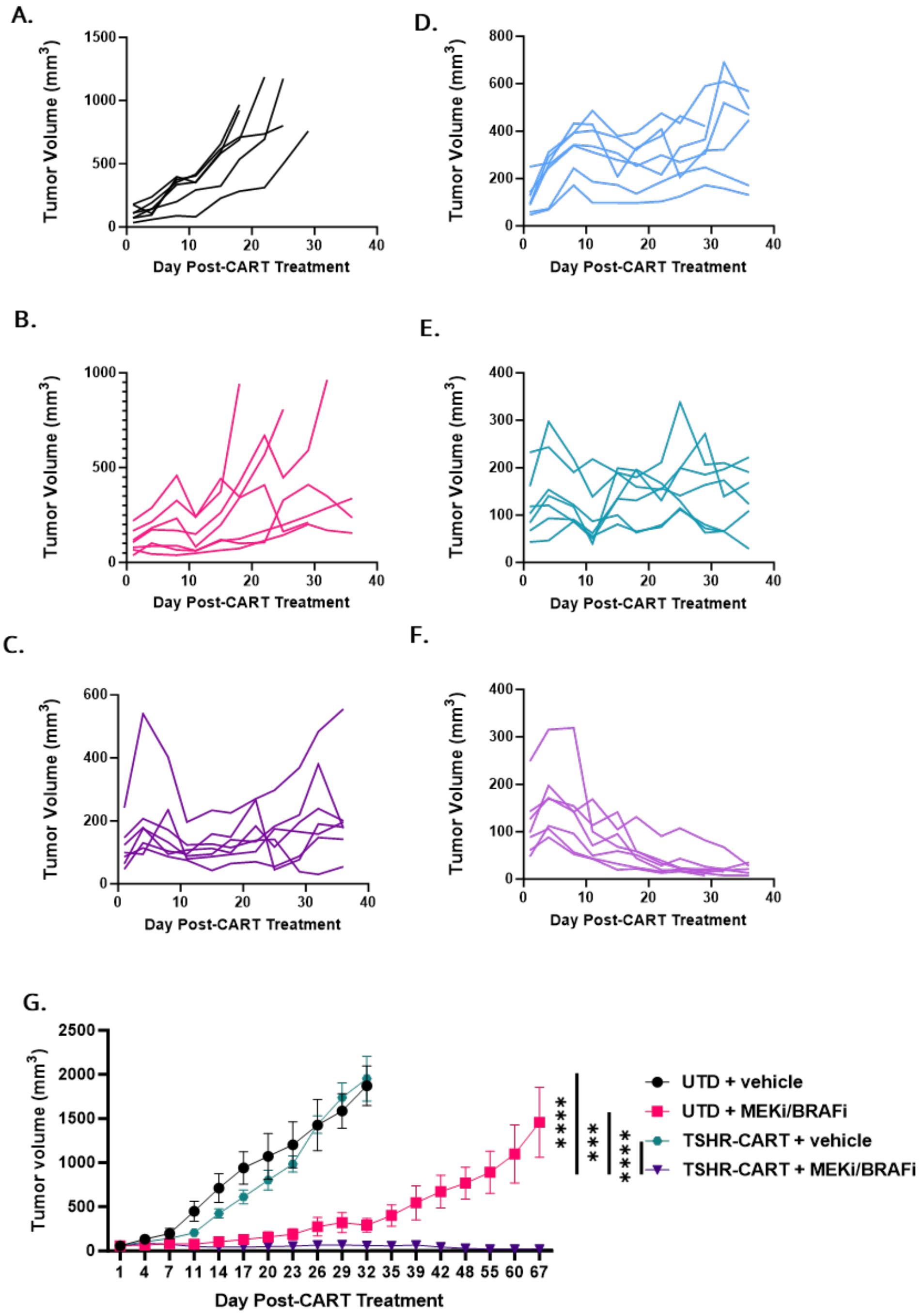


**Supplementary Figure S8. TSHR-CART cells showed the most potent antitumor activity when combined with concurrent MEKi/BRAFi administration. A-F)** Individual tumor growth curves are shown for each treatment group, including UTD + vehicle (**A**), sequential MEKi/BRAFi → UTD (**B**), concurrent MEKi/BRAFi + UTD (**C**), TSHR-CART + vehicle (**D**), sequential

MEKi/BRAFi → TSHR-CART (**E**), and concurrent MEKi/BRAFi + TSHR-CART (**F**). N = 7 mice/group. **G)** Tumor volume was assessed via serial caliper measurements (mean and SEM; ***p < 0.001, ****p < 0.0001, two-way ANOVA; n=7-15 mice per group).

Sup. Fig. S9

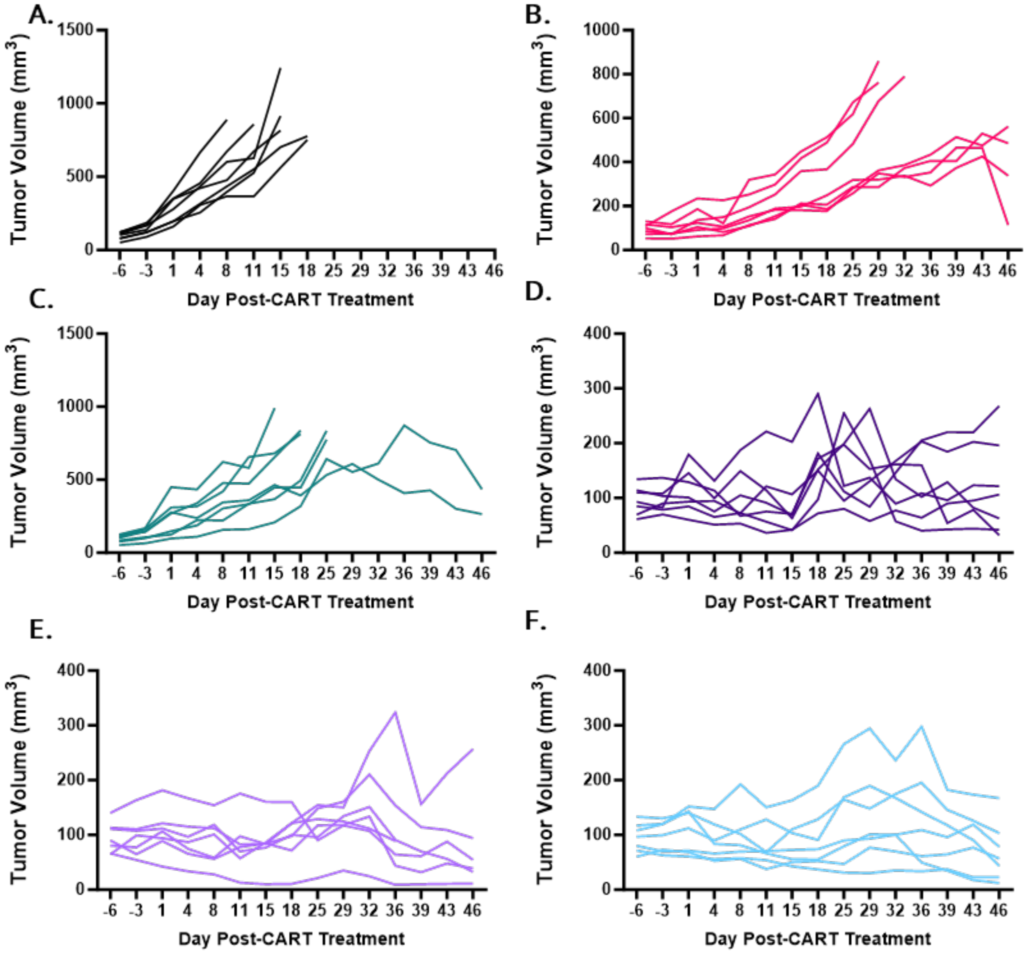


**Supplementary Figure S9. TSHR-CART cells showed the most potent antitumor activity when combined with MAPK inhibitor administration. A-F)** Individual tumor growth curves are shown for each treatment group, including UTD + vehicle (**A**), continuous MEKi/BRAFi + UTD (**B**), TSHR-CART + vehicle (**C**), MEKi/BRAFi (stopped after two weeks) + TSHR-CART (**D**), MEKi/BRAFi (stopped after 4 weeks) + TSHR-CART (**E**), and continuous MEKi/BRAFi + TSHR-CART (**F**). N = 7 mice/group.

**Supplementary Table S1: Detailed information of reagents used in this study.**

| Reagent | Company | Identifier |
|---|---|---|
| Goat anti-Human IgG Fc Recombinant Secondary Antibody (Alexa Fluor™ 647) | Invitrogen | A55749 |
| LIVE/DEAD™ Fixable Aqua Dead Cell Stain Kit, for 405nm excitation | Invitrogen | L34963 |
| APC anti-human CD45/clone 2D1 | BioLegend | 368511 |
| BV605 anti-human CD3/clone SK1 | BioLegend | 344835 |
| Pe-Cy7 anti-human CD4/clone SK3 | BD Biosciences | 557852 |
| BV421 anti-human CD8/clone RPA-T8 | BD Biosciences | 562428 |
| PE anti-human CCR7/clone G043H7 | BioLegend | 353203 |
| APC-Cy7 anti-human CD45RA/clone HI100 | BioLegend | 304151 |
| CellTrace™ CFSE Cell Proliferation Kit | Invitrogen | C34554 |
| Zombie R718™ Fixable Viability Kit | BioLegend | 423115 |
| APC-H7 anti-human CD3/clone SK7 | BD Biosciences | 560275 |
| MILLIPLEX® Human Cytokine/Chemokine/Growth Factor Panel A 38 Plex Premixed Magnetic Bead Panel | Millipore Sigma | HCYTA-60K-PX38 |
| FITC anti-human CD107a | BD Pharmigen | 560949 |
| Purified mouse anti-human CD28 | BD Biosciences | 348040 |
| Purified mouse anti-human CD49d | BD Biosciences | 340976 |
| Monensin Solution (1000x) | BioLegend | 420701 |
| CD3 Monoclonal Antibody (UCHT1), APC | Invitrogen | 17-0038-42 |
| PE-CF594 anti-human IL2/clone 5344.111 | BD Horizon | 562384 |
| BV421 anti-human GM-CSF/clone BVD2-21C11 | BD Horizon | 562930 |

| | | |
|---|---|---|
| PE-Cy7 anti-human MIP-1β/clone D21-1351 | BD Pharmigen | 560687 |
| BD FACS Lysing Solution 10X concentrate | BD Biosciences | 349202 |
| PE anti-human CD3/clone OKT3 | BioLegend | 317308 |
| BV421 anti-human CD45/clone HI30 | BioLegend | 304032 |
| APC-eFluor 780 anti-mouse CD45/clone 30-F11 | Invitrogen | 47-0451-82 |
| D-Luciferin, Potassium Salt | Gold Biotechnology | LUCK-100 |
| CD4 MicroBeads (Human) | Miltenyi Biotec | 130-097-048 |
| CD8 MicroBeads (Human) | Miltenyi Biotec | 130-045-201 |
| Dabrafenib (GSK2118436) | Selleck Chemicals | S2807 |
| Trametinib (GSK1120212) | Selleck Chemicals | S2673 |
| Anti-TSH Receptor/TSH-R antibody [EPR19751] | abcam | ab218108 |
| Anti-CD3 epsilon antibody [EP449E] | abcam | ab52959 |
| Goat anti-Human IgG Fc Recombinant Secondary Antibody (Alexa Fluor™ 647) | Invitrogen | A55749 |
| LIVE/DEAD™ Fixable Aqua Dead Cell Stain Kit, for 405 nm excitation | Invitrogen | L34963 |
| CountBright™ Absolute Counting Beads, for flow cytometry | Invitrogen | C36950 |
| BV605 anti-human CD3/clone SK1 | BioLegend | 344835 |
| Pe-Cy7 anti-human CD4/clone SK3 | BD Biosciences | 557852 |
| BV421 anti-human CD8/clone RPA-T8 | BD Biosciences | 562428 |
| PE anti-human CCR7/clone G043H7 | BioLegend | 353203 |
| APC-Cy7 anti-human CD45RA/clone HI100 | BioLegend | 304151 |
| CellTrace™ CFSE Cell Proliferation Kit | Invitrogen | C34554 |
| Zombie R718™ Fixable Viability Kit | BioLegend | 423115 |

| | | |
|---|---|---|
| APC-H7 anti-human CD3/clone SK7 | BD Biosciences | 560275 |
| MILLIPLEX® Human Cytokine/Chemokine/Growth Factor Panel A 38 Plex Premixed Magnetic Bead Panel | Millipore Sigma | HCYTA-60K-PX38 |
| FITC anti-human CD107a | BD Pharmigen | 560949 |
| Purified mouse anti-human CD28 | BD Biosciences | 348040 |
| Purified mouse anti-human CD49d | BD Biosciences | 340976 |
| Monensin Solution (1000x) | BioLegend | 420701 |
| FIX & PERM™ Cell Permeabilization Kit | Invitrogen | GAS004 |
| CD3 Monoclonal Antibody (UCHT1), APC | Invitrogen | 17-0038-42 |
| PE-CF594 anti-human IL2/clone 5344.111 | BD Horizon | 562384 |
| BV421 anti-human GM-CSF/clone BVD2-21C11 | BD Horizon | 562930 |
| PE-Cy7 anti-human MIP-1β/clone D21-1351 | BD Pharmigen | 560687 |
| APC-eFluor 780 anti-mouse CD45/clone 30-F11 | Invitrogen | 47-0451-82 |
| D-Luciferin, Potassium Salt | Gold Biotechnology | LUCK-100 |
| CD4 MicroBeads (Human) | Miltenyi Biotec | 130-097-048 |
| CD8 MicroBeads (Human) | Miltenyi Biotec | 130-045-201 |
| Dabrafenib (GSK2118436) | Selleck Chemicals | S2807 |
| Trametinib (GSK1120212) | Selleck Chemicals | S2673 |
| Anti-TSH Receptor/TSH-R antibody [EPR19751] | abcam | ab218108 |
| Anti-CD3 epsilon antibody [EP449E] | abcam | ab52959 |
